# Multiple omic investigations of freeze tolerance adaptation in the aquatic ectothermic vertebrate, the Amur sleeper

**DOI:** 10.1101/2022.05.10.491133

**Authors:** Wenqi Lv, Haifeng Jiang, Yuting Qian, Minghui Meng, Cheng Wang, Ning Sun, Yongrui Lu, Houhua Bing, Chengchi Fang, David M. Irwin, Shunping He, Liandong Yang

**Affiliations:** The Key Laboratory of Aquatic Biodiversity and Conservation of Chinese Academy of Sciences, Institute of Hydrobiology, Chinese Academy of Sciences, Wuhan 430072, China; College of Animal Science and Technology, Northwest A&F University, Yangling, Shaanxi 712100, China; Academy of Plateau Science and Sustainability, Qinghai Normal University, Xining 810016, P. R. China; State Key Laboratory of Genetic Resources and Evolution, Kunming Institute of Zoology, Chinese Academy of Sciences; University of Chinese Academy of Sciences, Beijing 100049, China; Diggers (Wuhan) Biotechnology Co., Ltd, Wuhan 430000, China; Center for Excellence in Animal Evolution and Genetics, Chinese Academy of Sciences, Kunming 650223, China; Department of Laboratory Medicine and Pathobiology, University of Toronto, Toronto M5S1A8, Canada; Banting and Best Diabetes Centre, University of Toronto, Toronto M5S1A8, Canada

**Keywords:** freeze tolerance, Amur sleeper, hypometabolism, cell stress response, cytoskeleton, cryoprotectant, nerve transmission

## Abstract

Freeze tolerance is an amazing overwintering strategy that enables ectotherms to occupy new niches and survive in cold climates. However, the genetic basis underpinning this ecologically relevant adaptation is largely unknown. Amur sleeper is the only known freeze-tolerant fish species that can overwinter with its entire body frozen in ice. Here, we sequenced the chromosome-level genome of the Amur sleeper and performed comparative genomic, transcriptomic, and metabolomic analyses to investigate this remarkable adaptation. Phylogenetic analyses showed that the Amur sleeper diverged from its close relative with no cold hardiness about 15.07 million years ago and revealed two unusual population expansions during the glacial epochs. Integrative omics data identified a synchronous regulation of genes and metabolites involved in hypometabolism and cellular stress response, and several related genes showed strong evidence of accelerated evolution and positive selection. Potential evolutionary innovations that might aid in freezing survival were found to be associated with the dynamic rearrangement of the cytoskeleton to maintain cell viability, redistribution of water and cryoprotectants to limit cell volume reduction, and inhibition in nerve activity to facilitate dormancy, demonstrating a coordinated evolution for this complex adaptation. Overall, our work provides valuable resources and opportunities to unveil the genetic basis of freeze tolerance adaptation in ectothermic vertebrates.

## Introduction

Freeze tolerance, the ability of an organism to withstand whole body freezing, is a striking winter survival strategy adapted by many ectotherms living in seasonally cold climates (Schmid 1982; Storey and Storey 1996b). At sub-zero temperatures, freeze-tolerant animals can withstand the conversion of as much as ~ 82% of their total body water into extracellular ice (Ramlov and Westh 1993). These animals may spend a prolonged state of frozen dormancy (days to months) in their hibernation states with the cessation of vital physiological functions including heartbeat, respiration, nerve conductance, and skeletal muscle movement, and return to active lives after thawing in the warm spring (Storey 1987, 1990). Although freeze tolerance has evolved multiple times across the animal kingdom ranging from insects, invertebrates, reptiles, and amphibians (Costanzo and Claussen 1990; Hans, et al. 1992; Loomis 1995; Layne Jr and Kefauver 1997; Bradley Shaffer, et al. 2013), it is actually a minority choice among vertebrates compared with common overwintering strategies like migration, hibernation and freeze avoidance (Costanzo and Lee 2013; Iwaya-Inoue, et al. 2018; Mohr, et al. 2020).

Most natural freeze tolerance refers to ice formation in extracellular spaces while resisting intercellular freezing to avoid damage of subcellular compartments and the cytoskeleton (Costanzo and Lee 2013; Storey and Storey 2017). Ice crystals exclude solutes greatly elevates the osmolality of extracellular fluids, producing a hyperosmotic stress that draws water out of cells causing them to shrink (Storey and Storey 2020). Besides physical and osmotic damage, freezing causes important consequences including hypoxia/anoxia, ischemia, dehydration and hypometabolism etc (Storey and Storey 2017; Toxopeus and Sinclair 2018). Moreover, reoxygenation, rehydration and reperfusion during thawing also accompany severe stresses (Giraud-Billoud, et al. 2019). To data, a suite of complex coordinated cellular, molecular, and physiological adaptations that confer freezing survival has been extensively and well explored in multiple hibernating reptile and amphibian species, especially the wood frogs (Zhang and Storey 2012; Bradley Shaffer, et al. 2013; Storey and Storey 2013, 2017; Costanzo 2019). These adaptations include mechanisms to manage extracellular ice volume and growth rate, strong metabolic rate depression coupling with selective activation of “survival” pathways to maintain stability, and accumulation of low-molecular-weight organic compounds as cryoprotectants. However, hitherto, the genetic basis of freeze tolerance in ectothermic vertebrates remains largely unknown.

The Amur sleeper, *Perccottus glenni* (Odontobutidae, Perciformes), is an aquatic species that can overwinter with its entire body frozen in ice, and probably the only freeze-tolerant vertebrate aside from reptiles and amphibians (Chai, et al. 2020). It is a limnophilic species native to the Amur River drainage in northeastern Asia, and has invaded European waters, leading to detrimental ecological impacts (Reshetnikov and Ficetola 2011; Xu, et al. 2014). Amur sleeper prefers small, stagnant waterbodies, which commonly freeze to the bottom in the winter. Before freezing, the Amur sleeper experiences long-term hypoxic or anoxic conditions in ice-covered waters until its whole body is encapsulated in ice (Reshetnikov 2003; Karanova 2009). Frozen dormancy can be maintained for up to three months (typically December to March), with revival occurring within a few hours of thawing (Chai, et al. 2020). This ecologically relevant freeze tolerance provides the Amur sleeper with a competitive advantage over other freshwater fishes that are unable to survive in such an extreme environment (yielding an avoidance of competitors and predators), allowing it to become one of the most widespread and successful fish invaders (Reshetnikov and Ficetola 2011). Therefore, the Amur sleeper could serve as a new model for investigations of adaptive freeze tolerance in ectothermic vertebrates, for which the majority of our knowledge comes from research in a few terrestrial amphibians. Moreover, the Amur sleeper has a small genome size and simple evolutionary history, thus providing an ideal opportunity to study the genetic basis of freeze tolerance in ectothermic vertebrates.

In this study, we generated a high-quality chromosome-level genome for Amur sleeper and a de novo reference genome assembly for *Neodontobutis hainanensis*, the closest relative to *P. glenni* with no cold hardiness (Lv, et al. 2020). First, we conducted comparative genomic analyses to explore the population histories and genetic changes of the Amur sleeper. Then we combined transcriptomic, and metabolomic analyses to better understand the molecular adaptations accompanying freeze tolerance. Our results not only gain insights into the genetic basis of this remarkable adaptation, but also provide useful genetic resources for future study.

## Results and discussion

### Genome Characteristics

We generated the first chromosome-level genome assembly for *P. glenii* using a combination of Nanopore long reads, BGISEQ-500 reads and Hi-C data (supplementary table S1). The genome size of *P. glenii* was estimated to be 827.25 Mb with a heterozygous ratio of 0.65% (supplementary fig. S1, supplementary table S2). Three assembly algorithms were used, and with the genome assembled by SmartDenovo finally selected based on continuity (supplementary table S3). After the removal of sequence redundancy, the genome size of the assembly was 710.22 Mb with contig N50 of 5.49 Mb (supplementary table S3). Further, we anchored and oriented 702 contigs (689.63Mb, ~97.09%) into 22 chromosomes (fig. 1A, supplementary fig. S2, supplementary table S4). Finally, a chromosome-level genome with contig and scaffold N50 equal to 2.96 Mb and 29.56 Mb, respectively, was generated (supplementary table S5). Over 99% of the short reads could be mapped to the genome, which covering 98.61% of the *P. glenii* genome assembly (supplementary table S6). Evaluation of the completeness based on BUSCO identified 91.2% complete and 3.3% fragmented genes (supplementary table S7). For comparative analyses, a de novo assembly of the *N. hainanensis* genome was performed and an individual with heterozygous ratio of 0.15% yielded a ~848 Mb assembly containing 8.221 contigs with the N50 of 1.34 Mb (supplementary table S5). A total of 97.21% short reads were mapped to the *N. hainanensis* assembly (supplementary table S6), and 93.10% complete BUSCO genes captured (supplementary table S7). The higher heterozygous rate in *P. glenii* may suggest an admixture of different clades during the its northwards expansion, and results in a reduction of genome size compared to *N. hainanensis* due to the removal of sequence redundancy.

**Fig. 1.**
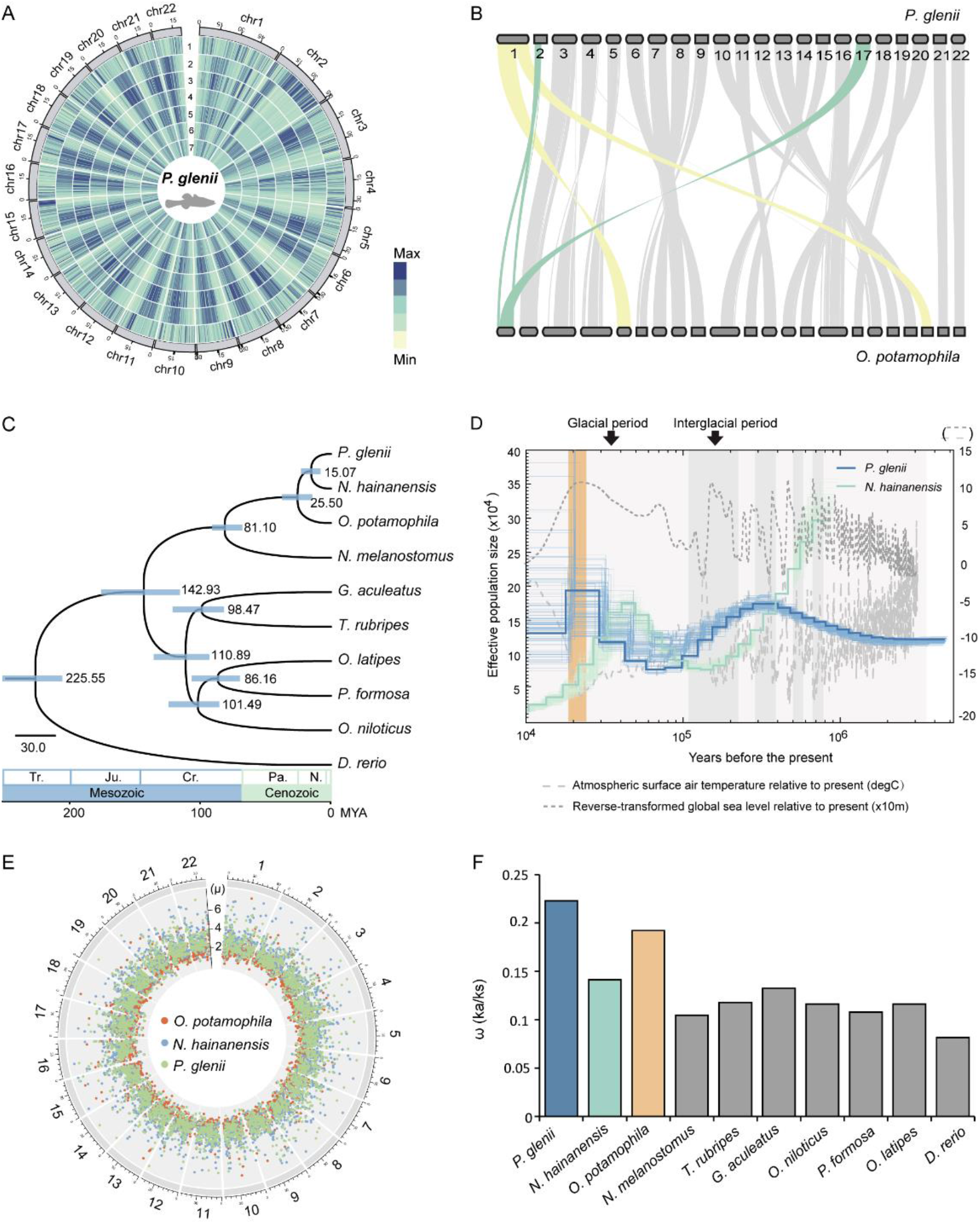
Evolutionary history of the Amur sleeper. (A) Circos plots showing the distributions of the genomic components in *P. glenii* with 500,000 bp windows. 1: Gene frequency, 2: Density of LINEs, 3: Density of LTRs, 4: Density of DNA, 5: Density of TRFs, 6: Density of SINEs, 7: Density of GC content. (B) Collinearity analysis of the *P. glenii* and *O. potamophila* genomes. (C) Phylogenetic tree and divergence times estimated for the Amur sleeper and nine other teleosts. Numbers near each node are the estimated divergence times, with the blue error bars indicating the 95% confidence levels. (D) Demographic history estimated by PSMC. Blue lines represent *P. glenii*, and green lines represent *N. hainanensis*. Orange frame represents the last glacial maximum (LGM). (E) Mutation rates for three species estimated across the genome. Number around the outside represent the chromosome ID for the *O. potamophila* genome. µ represents the mutation rate (×10^−9^ per site per year) of each window. (F) The ω(*Ka/Ks*) ratios of concatenated genes in ten species.

The GC content were 39.80% and 39.38 % for the *P. glenii* and *N. hainanensis* genome, respectively (fig. 1A, supplementary table S5). Approximately 48.13% and 45.99% of bases were identified as repetitive sequences in the two genomes by combining de novo and homology-based prediction methods (supplementary table S8). We predicted a total of 23,582 and 26,237 protein-coding genes in *P. glenii* and *N. hainanensis*, respectively, and approximately 97.45% of the protein-coding genes in *P. glenii* and 94.63% of those in *N. hainanensis* were successfully annotated using five public databases (supplementary table S9). We also compared the *P. glenii* genome karyotype with the genome of *O. potamophila*, a representative species also from Odontobutidae family. Only two chromosomal fission and fusion events were detected, indicating a conserved chromosomal evolution after divergence of the two species (fig. 1B). Taken together, these results revealed high-quality assembly and accuracy of annotation for the *P. glenii* and *N. hainanensis* genomes, which can be important genetic resources for further comparative and functional studies of freeze tolerance in ectotherms.

### Population history and Evolutionary rate

Phylogenetic analysis based on a set of 4,550 one-to-one orthologs from ten teleosts indicated that *P. glenii* was closest to *N. hainanensis* and form a monophyletic sister group to *Odontobutis* (fig. 1C and supplementary fig. S3). The reconstructed topology is consistent with topologies inferred by the mitochondrial genome and nuclear coding genes (Li, et al. 2018; Lv, et al. 2020). The divergence time between *P. glenii* and *N. hainanensis* in the present study was estimated at 15.07 Ma (7.94-23.02 Ma, 95% HPDs) in min-Miocene (fig. 1C).

Analysis using pairwise sequentially Markovian coalescent (PSMC) model (Li and Durbin 2011) revealed quite distinct demographic patterns for the two species (fig. 1D), which could be related to the significant climate oscillations involving glacial-interglacial cycles and sea level changes. Interestingly, two events of unusual population expansions were detected in the demographic history of Amur sleeper. The first expansion occurred before ~3 Ma and reached a peak ~ 0.3 Ma at the largest Quaternary glaciation (0.80–0.20 Ma) in the late Pleistocene, indicating that Amur sleeper has already developed a mechanism of cold adaptation at the Pliocene glaciation. This expansion corroborates well with a previous study that the Amur sleeper may have spread from the warm south to the cold north during Late Pliocene (2.58–3.60 Ma) (Li, et al. 2018). The subsequent declines coincided with the advent of the warm interglacial period, during which the rise in sea level caused by deglaciation resulted a dramatic reduction in freshwater habitats. Similarly, the second expansion occurred at ~ 70 Ka and reached a peak at the last glacial maximum (LGM, 26.5–19.0 ka, (Clark, et al. 2009)). In contrast, the population size of *N. hainanensis* dropped sharply since ~ 0.9 Ma and the only expansion occurred at ~ 0.15 Ma, followed by sharp declines predating the LGM. Overall, Amur sleeper maintains a relatively stable effective population size and has rich genetic diversity than *N. hainanensis*. This may be attributed to its strong resistance and adaptability, which can aid survival in extreme environments, thereby providing opportunities to expand into new ecological niches.

The development of freeze tolerance suggests adaptive evolution in the Amur sleeper, thus, we calculated its mutated and evolutionary rates. The mutation rate across the whole genome for the Amur sleeper is comparable to that of the other closely relative species (fig. 1E), however, our analyses revealed a higher evolutionary rate in the Amur sleeper, implying possible changes in the selection pressure experienced by *P. glenii* (fig. 1F, supplementary fig. S4).

### Transcriptomic and metabolic profiles over freeze/thaw

To understand the genetic regulatory mechanisms and metabolic adaptations, the transcriptome of the brain, liver, and muscle tissues and metabolomes of the liver and muscle tissues from three periods in a freeze/thaw episode, i.e., active autumn (AC), winter freezing (FR) (fig. 2A, supplementary movie S1), and early spring recovery (RE) (fig. 2B, supplementary movie S2), were analyzed for the Amur sleeper. Principal component analyses (PCA) of the transcriptomes showed that there were clear variations among the brain and liver tissues at different periods (supplementary fig. S5A). PCA analyses of the metabolomes showed large variations between the AC and FR and the AC and RE, while smaller differences between the FR and RE (supplementary fig. S5B). The number of differentially expressed genes (DEGs) and significantly different metabolites (SDMs) for the two comparisons i.e., AC vs FR and AC vs RE, were obviously more than those of FR vs RE (supplementary fig. S6A and B), indicating substantial changes in transcriptional and metabolomic profiles during the freezing and thawing periods. To better understand the potential functions of the regulation, GO and KEGG analyses of DEGs and SDMs were performed. Moreover, we further combined the results with comparative genomic analyses in order to provide clear insight into the genetic evolution and adaptive mechanisms of Amur sleeper’s freezing survival.

**Fig. 2.**
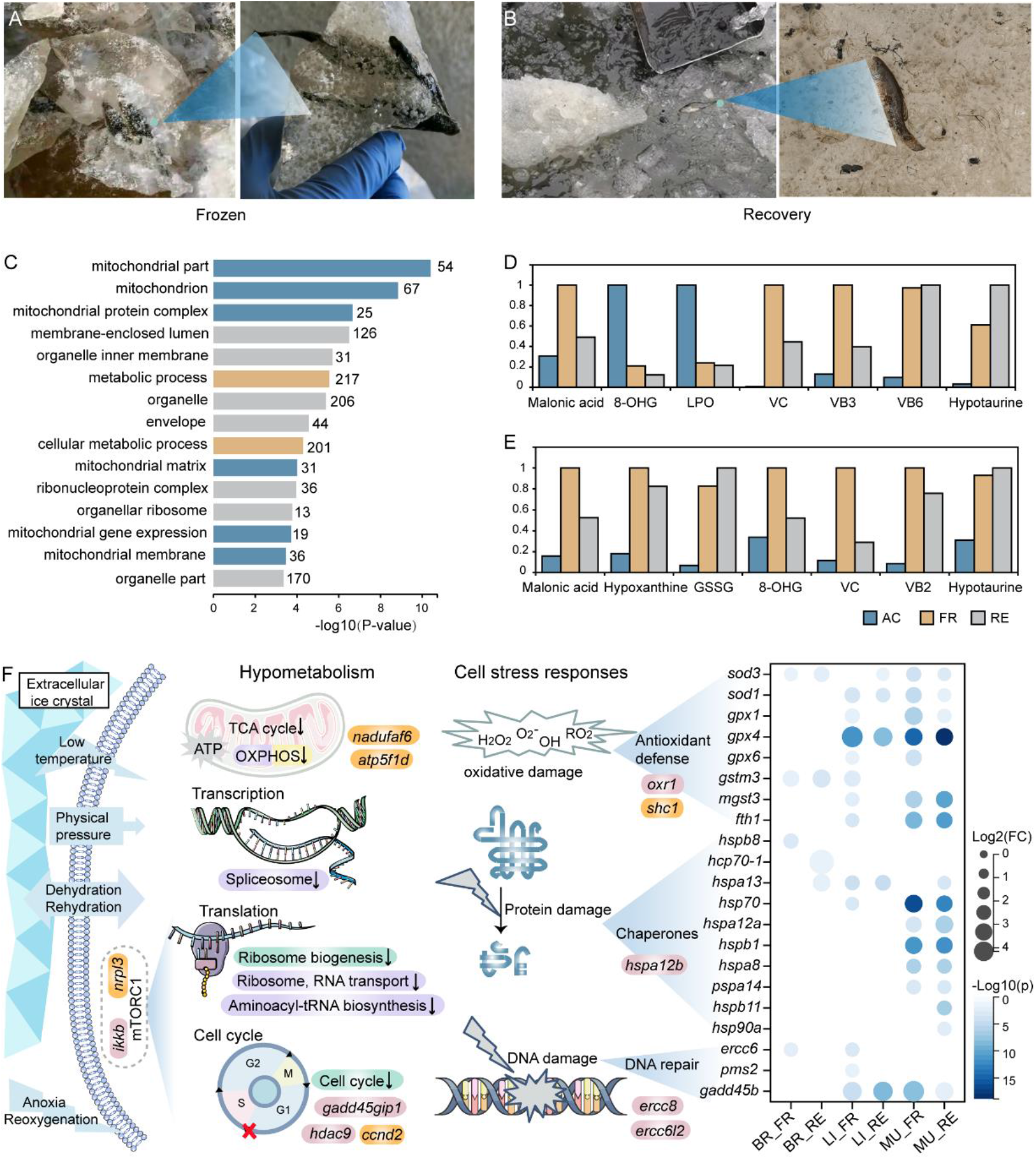
Genes and metabolites related to hypometabolism and cell stress response. (A and B) Illustrations of Amur sleeper during FR (A) and RE (B). (C) Top 15 significantly enriched GO terms for down-regulated genes in all three tissues at FR are shown. (D and E) Changes of metabolites related to metabolic inhibition and antioxidant defense in the liver (D) and muscle tissue (E). The relative levels of the metabolites are represented as bar graphs. Height of the column with the largest quantity is set as 100%, and that for the smaller quantities shown proportionally. (F) Pathways and genes associated with hypometabolism and cell stress responses. The significantly enriched pathways related to energy-expensive cell processes of down-regulated genes in brain (purple), liver (green), and muscle (yellow) tissues were shown. Genes are labeled with different colors to indicate positively selected genes (orange) and rapidly evolved genes (pink). The heatmap showed the transcriptional log2 (fold change) in expression of differently expressed genes in FR and RE relative to that in AC (BR: brain, LI: liver, MU: muscle, FC: fold change).

### Changes in genes and metabolites related to metabolic characteristics

Stress-induced metabolic rate depression (MRD) is the core adaptive strategy implemented by freeze tolerant animals to greatly decrease organismal energy demands, thereby permitting long-term survival using only their body fuel reserves (Carey, et al. 2003; Storey and Storey 2013). Here, transcriptomic profiles identified a set of 534 genes had significant changes in expression for all three tissues of Amur sleeper at FR (supplementary fig. 6A), among which, the downregulated genes were significantly enriched in the GO functional categories related to mitochondria (e.g., mitochondrial part, mitochondrion, and mitochondrial protein complex) (fig. 2C). The mitochondria produce most of the adenosine triphosphate (ATP) used by cells though mitochondrial energy metabolism, oxidative phosphorylation and tricarboxylic acid cycle reactions (Wu, et al. 2007). Importantly, *nadufaf6* and *atp5f1d*, which as components of the mitochondrial respiratory chain, were detected as positively selected genes (PSGs) (fig. 2F) in Amur sleeper. The observed signal of positive selection may have implications for regulating ATP synthesis. KEGG analysis revealed that down-regulated genes during FR in the brain and muscle tissue were significantly enriched in the oxidative phosphorylation pathway (fig. 2F, supplementary fig. S7A and C). Simultaneously, we found that malonic acid, an inhibitor of the tricarboxylic acid cycle by affecting succinate dehydrogenase (Lu, et al. 2018), increased about 3.28-fold and 6.34-fold in liver and muscle tissues during FR, respectively (fig. 2D and E, supplementary fig. S8A and B). Thus, the suppression of multiple mitochondrial pathways suggests a dramatic reduction in metabolic rate in the Amur sleeper during freezing. In addition, it is noteworthy that mitochondria are also the main cellular reactive oxygen species (ROS) generator, the inhibition would lead to a lower risk of ROS damage to macromolecules and organelles, including proteins, DNA, mitochondria, and cytoskeleton (Ou, et al. 2018).

The down-regulated genes were also enriched in pathways related to transcription, translation, and cell division (fig. 2F, supplementary Fig. S7A), which are all considered to be energy-expensive cell processes. Studies on wood frogs and western painted turtle demonstrated that cell cycle suppression is a general feature of freeze tolerance (Zhang and Storey 2012; Bradley Shaffer, et al. 2013). Remarkably, the *ccnd2* gene encodes G1/S specific cyclin D2 was positively selected in Amur sleeper and showed decreased expression at FR (fig. 2F). Inhibition of ccnd2 expression could arrest the cell cycle in G1 phase (Xiao, et al. 2021). *Gadd45gip1* and *hdac9* were identified as rapidly evolved genes (REGs), of which activity have been reported to be associated with cell cycle G1/S transition (Li and Durbin 2011). Moreover, we found a number of negative regulators of mTORC1 including *deptor, ddit4, tsc1* and *akt1* (Coronel, et al. 2022) significantly up-regulated at FR (supplementary fig. S9), indicating a strong depression of mTORC1 activity. Studies have reported that a reduction of mTORC1 activity could lead to subsequent inhibition of protein translation and cell cycle in hibernation animals (Logan, et al. 2019; Dias, et al. 2021). Furthermore, *nrpl3* and *ikkb* from mTOR signaling pathways were identified as PSG and REG, respectively (fig. 2F). Nrpl3 is a component of GATOR1, a complex involved in the inhibition of the mTORC1 (Baldassari, et al. 2016). Ikkb can activates the mTOR by mediating suppression of TSC1, a repressor of the mTOR pathway (Lee, et al. 2007). These genes that undergone marked genetic alterations may participle in the suppression of energy-expensive cell process in Amur sleeper during freezing, although their function still await further verification. Collectively, our results support a global MRD in Amur sleeper during frozen dormancy, which involves not only minimum energetic needs but also cellular defense strategy.

### Changes in genes and metabolites associated with cell preservation strategies

Both of freezing and thawing expose cells and organs to severe physiological stresses including anoxia/reoxygenation, dehydration/rehydration, ischemia/reperfusion, physical damage by ice and increased oxidative stress (Storey and Storey 2017; Zhang, et al. 2021). Therefore, an integrated suite of cellular stress response (CSR) to deal with these challenges is of paramount important to freeze-tolerant species. Core elements that constituted in the CSR including antioxidant defense, protein stabilization by chaperones, DNA damage repair, and so on (Kültz 2005). In metabolomic analysis, Vitamin C (Vc), as a well-known antioxidant was significantly increased 150.2-fold and 66.9-fold in the liver tissue during FR and RE, respectively, but only increased 8.7-fold and 2.5-fold respectively in the muscle tissue. Other vitamins, Vb3 and Vb6 have also shown antioxidant potential (Sinbad, et al. 2019), and showed consistent elevation at the two periods (fig. 2D and E, supplementary fig. S10 and11). Hypotaurine, which was evidenced as a strong antioxidant (Aruoma, et al. 1988), rose significantly in liver tissue at the FR (19.7-fold) and RE (29.7-fold), respectively, also with a slight increase in muscle tissue (3.0-fold and 3.22-fold, respectively) (fig. 2D and E, supplementary fig. S10 and11). Meanwhile, metabolites that can serve as markers of oxidative stress, such as glutathione oxidized (GSSG), hypoxanthine and 8-hydroxyguanosine (8-OHG) (Joanisse and Storey 1996; Hira, et al. 2014) showed notable increases in muscle tissue during FR and RE (fig. 2E, supplementary fig. S10). However, liver tissue had lower levels of 8-OHG and total lipid peroxidation (LPO) in these two periods, in agreement with its higher antioxidants amount (fig. 2D, supplementary fig. S11). Synchronously, our transcriptome profiles also identified a set of up-regulation genes that play crucial antioxidant roles during FR and RE. For example, the core antioxidant enzyme, superoxide dismutase (*sod1, sod3*), showed obviously higher expression levels in all three tissues, and glutathione peroxidases (*gpx1, gpx4*, and *gpx6*) were up-regulated in liver and muscle tissues. Other enzymes or proteins that have crucial antioxidant capacity, such as glutathione S-transferase (GST) isozymes (*gstm3, mgst3*) and ferritin (*fth1*) were also significantly increased (fig. 2F) (Tsuji, et al. 2000).

Heat shock proteins (HSPs) are the best-known chaperones that promote the refolding of denatured or misfolded proteins and prevent denaturation and aggregation of unfolded protein (King and MacRae 2015). Transcriptomic profiles revealed that the expression levels of number of HSP genes (including *hspb8, hsp70-1, hspa13, hsp70, hspa12a, hspb1, hspa8, hspa14, hspb11*, and *hsp90a*) significantly elevated especially in muscle during FR and RE (fig. 2F). Thus, the specific upregulated HSPs particular in muscle of Amur sleeper would contribute to stabilizing cellular proteome upon freeze/thaw. Moreover, genes involved in DNA damage repair, i.e., *ercc6, pms2*, and *gadd45b*, were also up-regulated during FR. Notably, some genes (*shc1, oxr1, hspa12b, ercc8* and *ercc6l2*) associated with CRS were found to be underwent positive selection and rapid evolution (fig. 2F). Shc1 and oxr1 has been reported to be important in protection from oxidative stress (Koch, et al. 2008; Sanada, et al. 2014). Ercc8 and ercc6l2 are essential factor in the transcription-coupled repair (TCR) pathway for DNA excision repair (Fousteri and Mullenders 2008; Tummala, et al. 2018). Overall, the synchronous regulation at transcriptional and metabolic levels and genetic changes provides strong evidence for the involvement of multiple cell preservation strategies, which would make important contributions to avoid metabolic damage and maintain cellular homeostasis in freeze tolerant species over freeze/thaw.

### Gene changes correlated with the cytoskeleton

The cytoskeleton consists of actin filaments (MF), microtubules (MT), and intermediate filaments (IF), not only helps control cell shape, bear external forces, and maintain the stability of internal cell structures, but also enables cells to carry out essential functions such as division and movement (Alberts, et al. 2002; Fletcher and Mullins 2010). During freezing, direct physical stress by extracellular ice and great changes in cell volume accompany freeze/thaw as well as destructive ROS would place significant consequences to the cytoskeleton. Importantly, multimeric cytoskeletal components quickly depolymerize at near-freezing temperatures, resulting in catastrophic functional impairment (Des Marteaux, et al. 2018; Ou, et al. 2018). Therefore, adaptive modifications of cytoskeletal related genes to cope with the challenges of cytoskeleton damage were likely necessary. As expected, our genomic comparisons of the Amur sleeper against other teleost revealed 230 gene families that had expanded significantly in size, and these families exhibited significant enrichments in functions associated with the cytoskeleton (e.g., cytoskeletal part, cytoskeleton organization, and microtubule-based process) (fig. 3A). Notably, three expanded gene families *(kntc2, spc24* and *haus3*) were related to kinetochore–microtubule attachment, a critical requirement for mitosis (fig. 3A, supplementary fig. S12). The *kntc2* and *spc24* gene families encode components of the NDC80 complex that is essential for stable kinetochore-microtubule anchoring and regulation of microtubules at the kinetochore (Umbreit, et al. 2012). Haus3, a component of the HAU augmin-like complex, has been reported to regulate the expression of the centrosome-related protein α-tubulin and the spindle-related protein γ-tubulin (Zhang, et al. 2019).

**Fig. 3.**
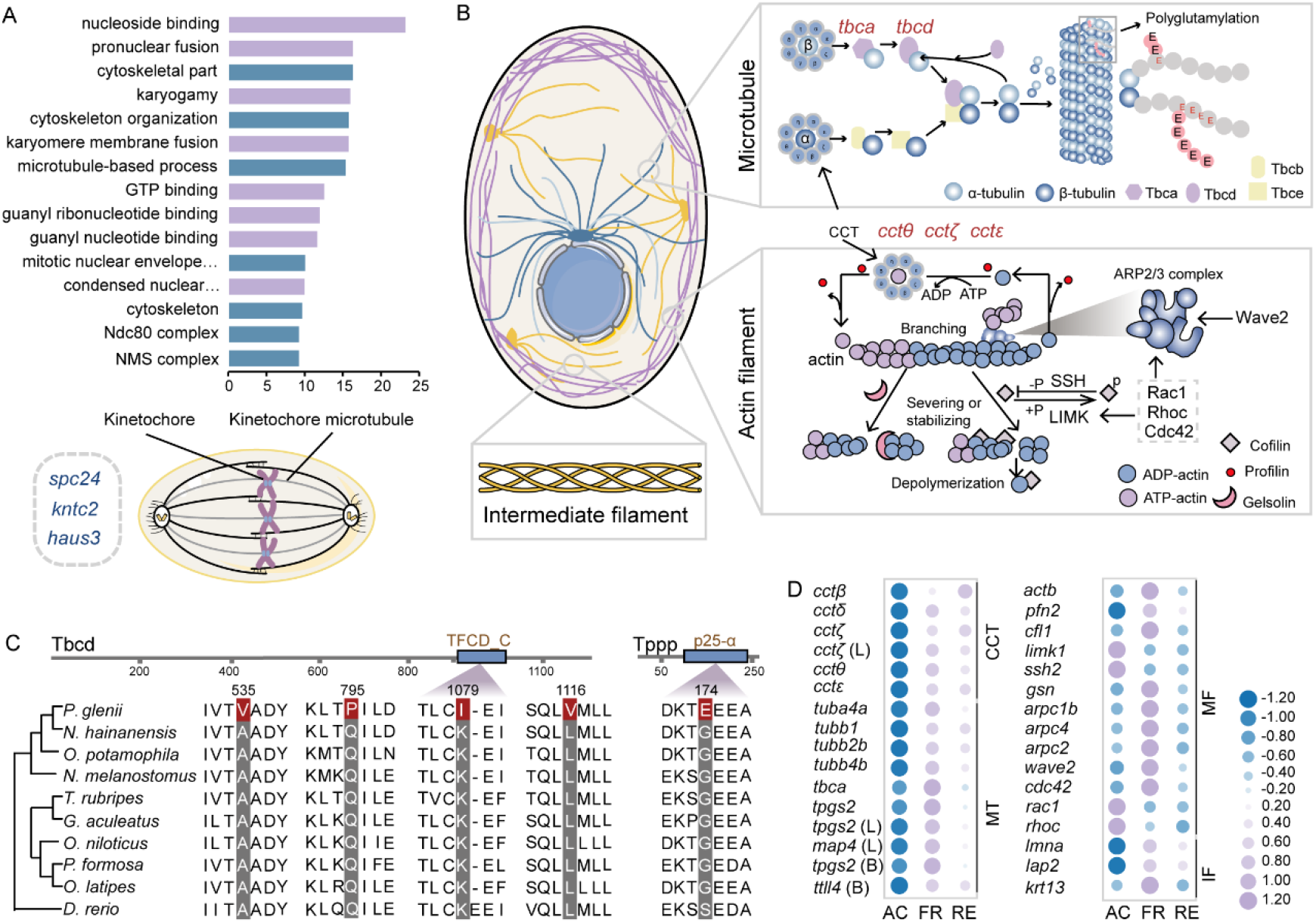
Changes in genes correlated with the cytoskeleton. (A) The top 15 significantly enriched GO terms for expanded gene families (up), and three expanded gene (down) that play crucial roles in mitosis (down) were shown. Blue columns represent cytoskeleton-related GO terms. (B) Schematic illustration of cytoskeletal composition, microtubule assembly, and regulation of actin cytoskeleton. Rapidly evolving genes are marked in red. (C) Sequence alignments for the positively selected genes, *tbcd* and *tppp*. Sites marked with red rectangles are positively selected sites. (D) Expression changes for genes related to the cytoskeleton. Purple represents higher expression levels, and blue represents lower expression levels. L and B represent liver and brain tissue, respectively (CCT: CCT complex; MT: microtubule; MF: actin filament; IF: intermediate filament).

Consistently, KEGG pathway enrichment analyses of REGs showed mostly significant enrichment in regulation of actin cytoskeleton and focal adhesin (supplementary fig. S13). Of particular interest, five chaperones *cctθ, cctζ, cctε, tbca*, and *tbcd*, exhibit significant evidence for elevated rates of evolution (fig. 3B). The genes *cctθ, cctζ* and *cctε* encode different subunits of chaperonin-containing t-complex polypeptide 1 (CCT complex) (Leitner, et al. 2012) are essential in the biogenesis of actin and tubulin to assemble MF and MT (Llorca, et al. 2001). Moreover, tbca and tbcd are tubulin-specific chaperones (TBCs) that bring together α- and β-tubulin subunits to form the assembly-competent heterodimer, which is required in the biogenesis of tubulin. Tbcd was further identified to be under positive selection, and showed four positively selected sites with one located in the TFCD-C domain (fig. 3C). Moreover, tubulin polymerization-promoting protein (*tppp*), a microtubule regulatory protein (Hlavanda, et al. 2002), was also identified as a PSG with a mutation (G174E) in p25-alpha domain (fig. 3C). Tppp not only promotes the incorporation of tubulin heterodimers into growing microtubule filaments (Hlavanda, et al. 2002), but also increases the level of microtubule acetylation, which is responsible for the stabilization of MT by inhibiting the activity of histone deacetylase 6 (Tőkési, et al. 2010). The genes display accelerated evolution in the Amur sleeper are pivotal in cytoskeleton stability, especially tubulin biogenesis, indicating a co-opted from cytoskeletal system to prevent irreparable structural damage and maintain normal function over freeze/thaw.

Our transcriptomic analyses lend further support to a dynamic cytoskeletal regulation in the Amur sleeper. Multiple genes involved in three components of the cytoskeleton exhibited significant expression fluctuations over FR and RE, especially in muscle tissue (fig. 3D). We observed significant upregulation of genes encoding subunits of CCT complex (*cctβ, cctδ, cctζ, cctθ*, and *cctε*) during the FR and RE stages (fig. 3D), suggesting the potential reassembly of dissociated cytoskeletal monomers. Indeed, the upregulation of the CCT complex involved in prevention of cold-induced actin depolymerization was also found in two insect species (Kayukawa and Ishikawa 2009; Zhang, et al. 2011). For microtubules, the genes *tubb1, tubb4b*, and *tubb2b* encoding β-tubulin, *tuba4a* encoding α-tubulin and chaperone gene *tbca* were up-regulated in the muscle during FR and RE with highest expression at FR (fig. 3D), likely indicative of maintenance of tubulin pool, which becomes more important at low temperatures that give rise to MT destabilization. Meanwhile, genes involved in post-translational modification (PTM) of tubulins (*map4, tpgs2* and *ttll4*) (Wang, et al. 2022) also exhibited a tissue-specific elevation at FR (fig. 3D). Such alterations may be important in promoting MT cold stability. With respect to more cold-resistant MF, we observed a highly expressed *actb* with a set of genes encoding actin-binding proteins profilin (*pfn2*), gelsolin (*gsn*), cofilin (*cf1*), and arp2/3 complex (*arpc1b, arpc4* and *arpc2*) that required for the polymerization and depolymerization of actin filaments (Pollard 2016) showed significant up-regulation in muscle tissue during FR. Simultaneously, upstream regulators of cofilin and arp2/3 complex such as *limk1, ssh2, wave*2, *cdc42, rac1*, and *rhoc* (Campellone and Welch 2010; Mizuno 2013) also exhibited obviously different expression patterns, representing a dynamic regulation on actin cytoskeleton (fig. 3B and D). Furthermore, we also found that genes e.g., *krt13, lmna*, and *lap2* that participate in assembly or PTM of IF (Naetar, et al. 2017; Wu, et al. 2017) showed significant expression fluctuation in muscle tissue during FR and RE (fig. 3D). Real-time quantitative PCR (qRT-PCR) analysis of selected genes were consistent with the transcriptome analysis (supplementary fig. S14A and B). Collectively, our analyses provide gene evolution and expression evidence for adaptive modification of cytoskeleton, and thus may play vital roles in maintaining cell viability after enduring freeze/thaw.

### Cryoprotectants and transmembrane transporters

A well know strategy for freezing survival is the accumulation of large quantities of low molecular weight cryoprotectants not only reducing ice formation but also limiting cell shrinkage via osmic effects (Storey and Storey 1986, 1996b). Sugars (e.g. glucose, trehalose), polyhydric alcohols (e.g. glycerol, sorbitol) and amino acid (e.g. proline) are common colligative (concentration dependent) cryoprotectants among cold-hardy amphibians and insects, and urea contributes in some cases in amphibians (Storey and Storey 2017; Toxopeus and Sinclair 2018). Our metabolomic profiles identified a variety of putative cryoprotectants showed distinct fluctuations in muscle and liver of Amur sleeper during FR and RE. Unexpectedly, glucose was the top among sugars, but only showed an apparent increase (3.6-fold) in muscle during FR (fig. 4A). Myo-inositol and glycerol were found to be the highest polyhydric alcohols in both tissues but with no significant fluctuations during FR while the followed arabitol had a remarkable rise in muscle (5.8-fold) (fig. 4A). Despite a global repression in amino acid metabolism, aspartic acid showed a substantial augment (7.88-fold) in muscle (fig. 4A), suggesting a potential protective role. This higher accumulation but less variation in contents for potential cryoprotectants in Amur sleeper likely indicative of a seasonal acquisition but not a freeze response, which agree with the contradictory reports regarding the changes of glucose or glycerol levels in different freeze tolerant frogs (Layne Jr and Jones 2001; Irwin and Lee 2003; Niu, et al. 2018). On the other hand, a recent study has suggested that multiple cryoprotectants (myo-inositol, proline, and trehalose) contribute to freeze tolerance largely via non-colligative mechanisms with each molecular is interchangeable and has a unique cryoprotective function (Toxopeus, et al. 2019).

**Fig. 4.**
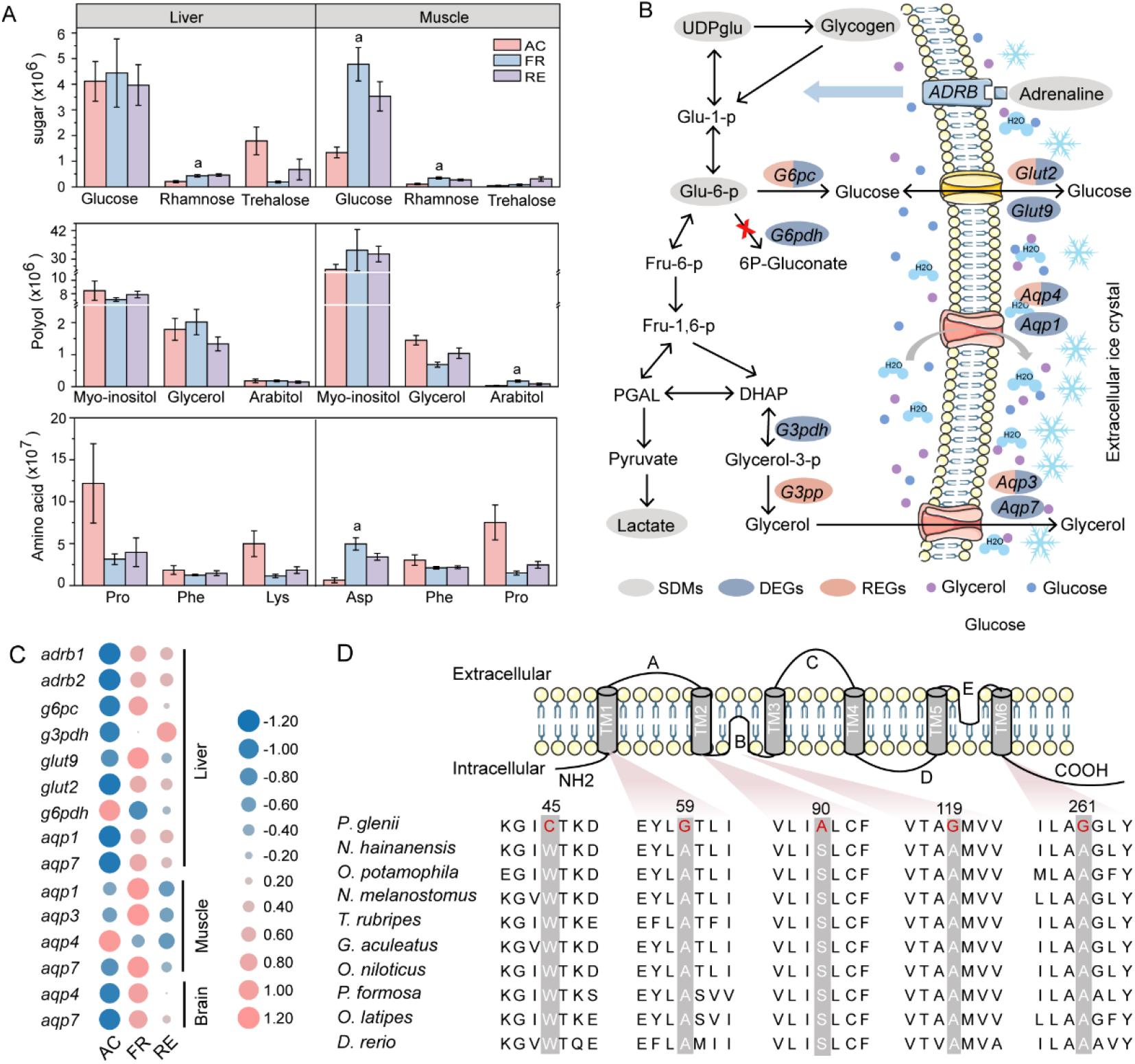
Putative cryoprotectants and their movement with water. (A) The top three metabolism with the highest content in sugar, polyhydric alcohols (polyol), and amino acid, respectively. The letter a indicates that the content increases significantly during FR stage. Error bars represent the mean ± S.D. (n = 6). (B) Schematic depiction of biosynthetic pathways for glucose and glycerol, and movement of water and cryoprotectants. (C) Expression changes of genes related to cryoprotectant. Pink represents higher expression levels, and blue represents lower expression levels. (D) Multiple-sequence alignments of *aqp4* amino acid sequences of Amur sleeper and other nine teleosts, and the specific mutations are marked in red.

During freezing, glycogenolysis is clearly the dominant pathway leading to produce cryoprotectants glucose and polyhydric alcohols. In Amur sleeper liver, we observed a sharp drop in glycogen level (as much as about 133-fold) and obvious rise in UDP-glucose (6.40-fold) and glucose-6-phosphate (2.92-fold) (fig. 4B, supplementary fig. S15). Meanwhile, the significant lactate accumulation at FR indicates glucose served as fuel for anaerobic metabolism (supplementary fig. S15). β-adrenergic signaling (the fight-or-flight response) response triggered by adrenaline has been linked with the rapid activation of glycogenolysis (Storey and Storey 1996a). Consistently, adrenaline rose 2.20-fold and the expression of beta-1 adrenergic receptor (*adrb1*) and beta-2 adrenergic receptor (*adrb2*) were significantly up-regulated (supplementary fig. S15, fig. 4B and C) at FR. Moreover, glucose-6-phosphatase (*g6pc*), which convert Glu-6-p to glucose in the terminal step of the synthesis of glucose (Foster, et al. 1997), was found to be significantly up-regulated (fig. 4B and C). Simultaneously, glucose 6-phosphatedehydrogenase (*g6pdh*) decreased obviously, this suppression could prevent Glu-6-p from being used for other purposes (Cowan and Storey 2001). It is notable that *g6pc* of the Amur sleeper was also identified as a REG and has nine specific amino acid replacements (fig. 4B, supplementary fig. S16A), four of them located in transmembrane domains (supplementary fig. S16B). Moreover, glycerol-3-phosphate phosphatase (*g3pp*), a crucial enzyme for glycerol biosynthesis (Raymond 2015) experienced an elevated rate of evolution (fig. 4B). These genetic changes might promote glucose and glycerol synthesis in the liver, which could be exported and circulated throughout the entire body for cryoprotection before freezing.

The formation of ice crystals and the accumulation of cryoprotectants have profound effects on the water content of cells as well as elevating osmolality (Storey and Storey 2013). The Aquaporins (AQPs) play an essential role in rapid osmoregulation as they allow for the facilitated diffusion of water and osmolytes across cell membranes (Hill, et al. 2004). There are two subsets of the AQP family, AQP1, AQP2, AQP4, and AQP5 that are permeated only by water and AQP3, AQP7 and AQP9 that can also be permeated by glycerol and even larger solutes (Verkman and Mitra 2000). In this study, the *aqp3* and *aqp4* were identified as REGs in the Amur sleeper (fig. 4B), and *aqp4* has five specific amino acid mutations that can cause polarity changes. These mutations including three alanine (A) to glycine (G) substitution occurred at highly conserved sites in the corresponding proteins of other fish species (fig. 4D). Furthermore, the generally increased expression levels of *aqp1, aqp3, aqp4* and *aqp7* in different tissues at FR provided functional evidence that these AQPs are extremely important to control ice content, cell volume, and maintain fluid homeostasis in Amur sleeper (fig. 4C). For glucose transport, transcript levels for members of the glucose transporter (GLUT) family glut2 and glut9 were significantly increased in liver at FR in Amur sleeper (fig. 4B and C), suggesting a rapid uptake of cryoprotective glucose during the onset of freezing. Moreover, glut2 was found to be underwent rapid evolve in Amur sleeper and up-regulated during RE (fig. 4B). Indeed, glut2 is a unique bidirectional transporter also allows for hepatocytic reuptake of glucose during thawing to restore glycogen pools and mitigate hyperglycemia (Storey and Storey 1986; Mueckler and Thorens 2013). Such dynamic regulation of glut2 expression is crucial to surviving freezing and thawing in organisms that employ glucose as cryoprotectant, which has been demonstrated in amphibians (Storey and Storey 1988; Rosendale, et al. 2014). The expression trends of genes that were found to be regulated in this part were validated by qRT-PCR analysis (supplementary fig. S14D and E). Taken together, genes associated with metabolic enzymes and transmembrane transporters may contribute to freeze tolerance by facilitating cryoprotectants synthesis and redistribution of water and cryoprotectants, although the effects of these mutations still need further investigation.

### Gene changes correlate with nerve activity

Freeze tolerant animals endure a prolonged state of frozen dormancy with interrupted nerve transmission (Storey and Storey 2017). Thus, a mechanism involved in the entry/exit and maintenance of dormant state over freeze/thaw is likely a key innovation, but this subject has received almost no attention in ectothermic vertebrates. It has been demonstrated that suppression of the central nervous system (CNS) plays a key role in the entrance into hibernation in mammals (Tamura, et al. 2005; Mohr, et al. 2020). Therefore, depressed neurotransmission activity in freeze tolerant ectothermic vertebrates when facing freezing would be expected. In the present study, we found a total of nine copies of the adenosine A1-receptor (*adora*) in the *P. glenii* genome, seven of which are tandemly duplicated and situated between *slc12a5* and *cyp24a1* genes (fig. 5A). This receptor is generally linked to the inhibition of the release of neurotransmitters with its most prominent inhibitory action on the excitatory glutamatergic system (Dunwiddie and Masino 2001). Such significant expansion might be important for the induction or maintenance of the dormant state in the Amur sleeper by suppressing neurotransmission. Interestingly, we found that *gabrg2*, encoding the γ2 subunit of gamma-aminobutyric acid receptor type A that mediates the inhibition of the CNS by combining with gamma-aminobutyric acid (GABA), possesses a strong signal of positive selection in the Amur sleeper. Remarkably, a total of 14 positively selected sites were detected, which is also highly conserved in human, mouse, and chicken (fig. 5B). Among them, 12 were located in the extracellular ligand-binding domain (fig. 5B). Furthermore, our transcriptomic analyses identified multiple genes (*gad2, glna, snat2, snat3, pkaca* and *pkcb*) in GABAergic synaptic pathway exhibited increased expression in brain tissue during FR and RE. Exposure of zebrafish to GABA-enhancing drugs, and mice fed with high GABA–containing black sicky rice giant embryo have antianxiety effects (Stewart, et al. 2011; Jung, et al. 2017). Thus, natural selection on gabrg2 and increased expression of the genes involved in GABAergic synaptic might have important antianxiety role, which perhaps could relieve freezing-induced stresses and facilitate entering dormancy.

**Fig. 5.**
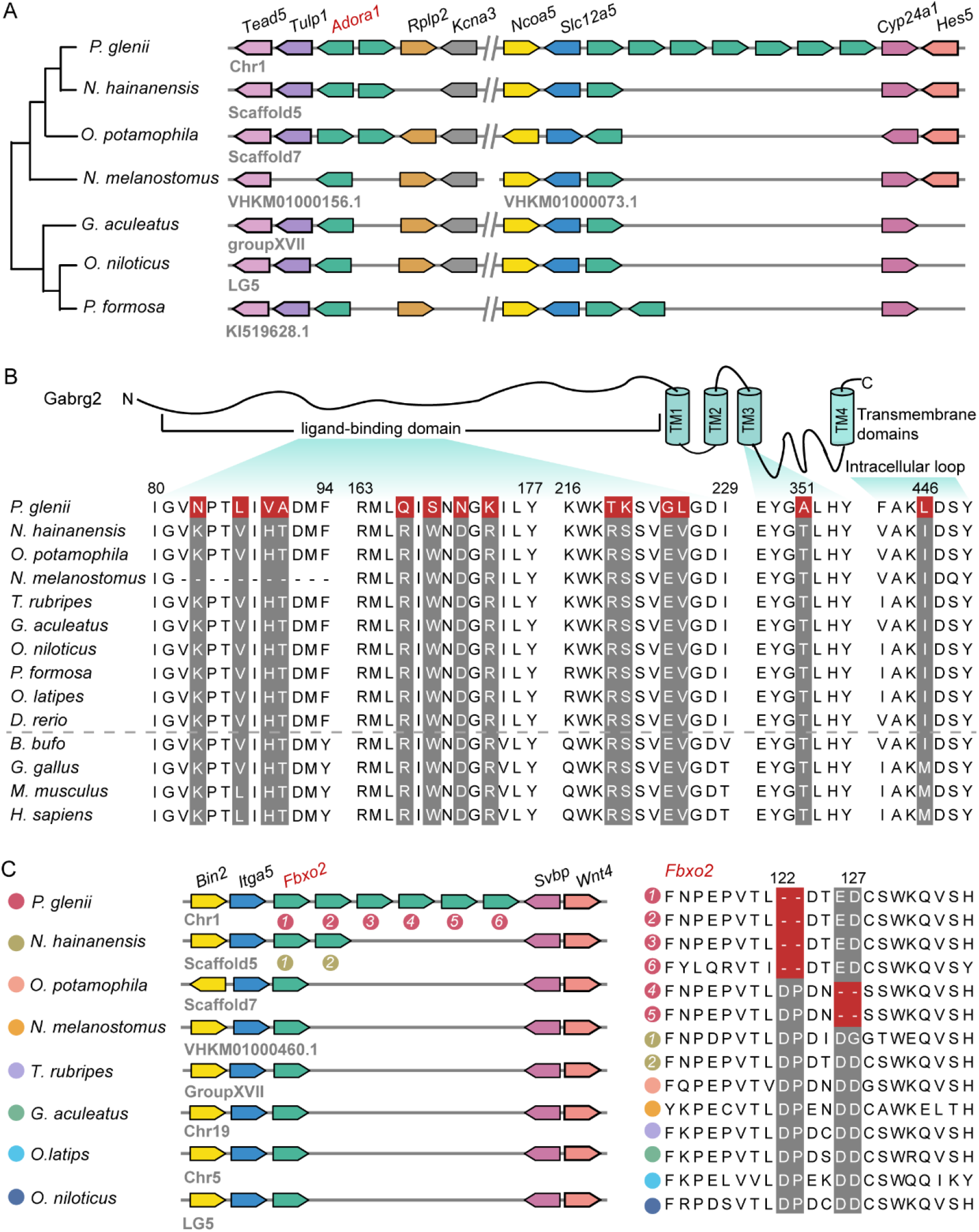
Expanded and positively selected genes related to nerve activity. (A) Expansion of the adenosine A1-receptor (*adora*) gene family. Seven copies of the Amur sleeper *adora* gene are arranged in tandem. (B) Schematic representation of gabrg2 protein, and the sequence alignment of Amur sleeper gabrg2 with nine teleosts and four other vertebrates (*B. bufo*: frog, *G. gallus*: chicken, *M. musculus*: mouse, *H. sapiens*: human).14 positively selected sites in Amur sleeper are boxed in red. (C) F-box only protein 2 (*fbxo2*) in the Amur sleeper is tandemly duplicated (left). Alignment of six copies of fbxo2 in Amur sleeper with other teleosts show specific deletions of 2-amino acid (right).

In concert with the receptors that mediate neural inhibition, we found metabotropic glutamate receptor 5 (*mglur5*) that modulates cell excitability and synaptic transmission, was under positive selection in the Amur sleeper. Two positively selected sites (C607L and L608T) were identified on the conserved dimer interface (supplementary fig. S18). The mglur5 receptor exists as dimers and monomers only contact via the extracellular domains in the inactive state, however, the dimer will undergo massive conformational change that bring the seven transmembrane domains closer together and into contact when it is activated (Llinas del Torrent, et al. 2019). In addition to increasing neuronal excitability, mglur5 also play important roles in the induction of long-lasting forms of synaptic plasticity, long-term depression (Niswender and Conn 2010). These two mutations may change the nerve conductance in the Amur sleeper and thus may have implications for its neural activity maintenance while frozen and reactivation upon thawing.

Besides genetic changes related to neurotransmission, ubiquitination (Ub) represent a dynamic PTM that precisely modulates the functional neuronal circuits. In Amur sleeper genome, we found six copies of f-box only protein 2 (fbxo2) that were tandemly duplicated and they situated between *bin2* and *svbp*. Particularly, two Amur sleeper-specific deletions of amino acid that could affect the three-dimensional structure of the protein were detected in all copies of these proteins, which was validated by mapping to both full-length and NGS transcripts. (fig. 5C, supplementary fig. S19). Fbxo2 is a subunit of the ubiquitin protein ligase complex SCF, which function in Ub-mediated degradation of GluN1 subunit of the NMDA receptors that mainly mediate neural excitatory transmission (Otsu, et al. 2019). GluN1 knockout mice have hyperactivity compared to wild-type mice (Segev, et al. 2020). Given that fbxo2 has been shown to facilitate the degradation of the GluN1 (Atkin, et al. 2015), the genetic changes of fbxo2 in Amur sleeper may suggest an inhibition of excitatory transmission. Taken together, the Amur sleeper-specific genetic innovations might thus indicate a role for adaptive evolution in function of the nervous system. Analysis of the variations in these genes may yield insights into how nerve regulation is coordinated upon freezing and thawing. Additionally, genes reported here are associated with multiple neurological and neuropsychiatric disorders in human, we expected that the genetic changes in the Amur sleeper may provide useful information for studies in mental illnesses and medical anesthesia.

### Conclusion

Freeze tolerance, a fascinating example of complex animal adaptations, has been extensively investigated in multiple hibernating reptile and amphibian species. However, the genetic basis for freezing survival remains unclear and the underlying molecular mechanisms are still inadequate understood in ectothermic vertebrates. Our study demonstrates the strengths of multi-omic methods to shed light on this adaptation in the only known fish species with freeze tolerant ability. Using an integrated analysis of multi-omic data, we revealed a suite of coordinated molecular adaptations in the Amur sleeper related to hypometabolism and cell repair that may mitigate the detrimental effects of freezing and thawing. Many significant genetics changes correlated with cytoskeletal stability, osmotic regulation, and nerve activity could be regarded as evolutionary innovations, which lay a blueprint for further functional characterization. This study not only provides useful genomic resources and insights into freeze-tolerant adaptation in ectothermic vertebrates, but also has potential implications for the development of better cryopreservation technologies and the unveiling of the causes of mental diseases in biomedical field.

## Materials and Methods

### Genome sequencing and de novo assembly

Wild individuals of *P. glenii* and *N. hainanensis* were collected from Heilongjiang and Guangxi Province, respectively. All experiments in this study were approved by the Institutional Animal Care and Use Committee of Institute of Hydrobiology, Chinese Academy of Sciences (Approval ID: Y21304506), and conducted in compliance with the relevant guidelines. Muscle tissues from *P. glenii* and *N. hainanensis* were used for DNA extraction and genome sequencing. For *P. glenii*, a total of 70.29 Gbp of long reads and 101.02 Gbp of short reads were generated. For *N*.*hainanensis*, a total of 86.06 Gbp of long reads and 121.71 Gbp of short reads were generated. The long reads were sequenced using the PrometlON DNA sequencer on the Oxford Nanopore platform and the short reads were generated on the BGISEQ-500 platform. For Hi-C sequencing of *P. glenii*, liver tissue was used for the extraction of DNA for library preparation. Hi-C libraries were then sequenced on the Illumina NovaSeq platform, and a total of 95.60 Gbp Hi-C reads were generated (supplementary table S1).

We adapted the KmerFreq_AR program from SOAPdenovo2 package, which is based on k-mer distribution, to estimate the genome size with about 45 Gb of BGISEQ-500 short reads filtered using fastp (Chen, et al. 2018). The estimated genome size of *P. glenii* and *N. hainanensis* were 827.25 Mb and 840.86 Mb, respectively (supplementary table S9). First, Nanopore long reads of the two species were corrected using the NextCorrect modules of NextDenovo (https://github.com/Nextomics/NextDenovo). For *P. glenii*, we used wtdbg2 (Ruan and Li 2020), Flye (Kolmogorov, et al. 2019), and Smartdenovo (https://github.com/ruanjue/smartdenovo) for de novo assembly with corrected long reads due to the high heterozygosity of the estimated genome characteristics (supplementary fig. S1, supplementary table S9). Next, we applied three rounds of polishing using filtered short reads with Pilon1.23 (Walker, et al. 2014). We also filtered the redundant contigs caused by high heterozygosity using the script fasta2homozygous.py from Redundans (https://github.com/lpryszcz/redundans). The quality of these three genomes were assessed. Finally, Hi-C reads were aligned to the best assembly version via Bowtie 1.2.2 (Langmead 2010). We then used Juicer v1.5 (Durand, Shamim, et al. 2016) and 3D-DNA (Dudchenko, et al. 2017) to anchor the draft genome onto 22 chromosomes. Juicerbox Assembly Tools (Durand, Robinson, et al. 2016) was used to visualize and improve the assembly quality. For *N. hainanensis*, the Nanopore reads were assembled by wtdbg2 and then three rounds of polishing using NSG short reads were applied. To estimate the quality of the two genomes, short reads were mapped back to the genome using BWA-MEM (Li and Durbin 2009). Completeness of the two genomes was evaluated using Busco v3 (Simão, et al. 2015) with the actinopterygii_odb9 database.

### Genome prediction annotation

We combined RepeatMasker v4.06 (Tarailo - Graovac and Chen 2009) with RepeatProteinMask v4.06 for homology repeat sequence prediction by aligning the genome sequences against the RepBase library. For de novo repeat prediction, we adopted RepeatModeler v1.08 along with LTR-FINDER v1.06 (Xu and Wang 2007) based on the de novo repeat library.

We used three different methods, namely, ab initio annotation, homology annotation and transcriptome-based annotation, to predict the whole gene set for the two genomes. Briefly, Augustus (Stanke, et al. 2008), Snap (Leskovec and Sosič 2016) and GeneScan (Burge and Karlin 1997) were used for de novo gene prediction based on the repeat masked genome sequences. The Augustus and Snap programs were trained with the transcript and zebrafish genome training set, respectively. For homology-based annotation, protein sequences from *Oreochromis niloticus, Oryzias latipes, Danio rerio, Anabas testudineus, Neogobius melanostomus* and *Poecilla formosa* were downloaded from Ensembl and aligned to the two genomes using the TBLASTN program. GeneWise (Birney, et al. 2004) was used to identify accurate gene structures for the alignment produced by TblastN. In addition, GeMoMa (Keilwagen, et al. 2019) was used for homology-based prediction with the zebrafish genome as a reference genome. For transcriptome-based annotation, a total of 62.49 Gb of RNA-seq reads from ten tissues of *P. glenii* and 87.30 Gb from nine tissues for *N. hainanensis* were generated by BGISEQ-500 (supplementary table S10). We also performed full-length transcriptome sequence, a total of 8,570,679 subreads with a mean length of 2,009 bp for *P. glenii* and 16,621,922 subreads with a mean length of 1719 bp for *N. hainanensis* were generated in PACBIO_SMRT platform (supplementary table S11). RNA-seq reads were mapped to the genomes using Hisat2 (Kim, et al. 2015), and then transcripts were generated using StringTie (Pertea, et al. 2015). Full-length transcriptome data were used to construct consensus sequences using IsoSeq3 (https://github.com/PacificBiosciences/IsoSeq), and subsequently mapped to the genomes with Gmap (Wu and Watanabe 2005). Both types of transcripts were then processed with PASA (Haas, et al. 2003) to obtain final results. Finally, all gene models generated from these three strategies were integrated with EVidenceModeler (EVM) (Haas, et al. 2008). Functional annotations of the predicted gene set were obtained by mapping to public functional databases, including SwissProt, NCBI-Nr, KEGG, GO and InterPro.

### Syntenic relationship with the dark sleeper genome

To evaluate the consistency of the Amur sleeper with its close relative, the dark sleeper (*Odontobutis potamophila*), we assembled and annotated the genome of dark sleeper based on raw data downloaded from NCBI (Jia, et al. 2021). First, the coding sequences from 22 autosomal chromosomes of the two species were aligned by LAST v942 (Kielbasa, et al. 2011). The results were then subjected to MCscan (Tang, et al. 2008) to identify syntenic blocks.

### Phylogenetic analysis

In addition to *P. glenii* and *N. hainanensis*, genomes of eight other teleosts, including *O. potamophila, O. niloticus, O. latipes, D. rerio, N. melanostomus, P. Formosa, Gasterosteus aculeatus*, and *Takifugu rubripes* were used to perform comparative genomic analyses. First, we identified orthologous gene clusters using OrthoFinder v2.3.4 (Emms and Kelly 2015) with default parameters. A total of 4550 single-copy genes were identified in the 10 species. Protein sequences for each single-copy orthologue were aligned using MAFFT v7.310 (Katoh and Standley 2013) and then corresponding coding sequence alignments were obtained with pal2nal v14 (Suyama, et al. 2006). We removed poorly aligned regions of each CDS alignment using Gblocks v0.91b (Talavera and Castresana 2007) with the codon model. Then the alignments of less than 50 codons were discarded (Wu, et al. 2021). All the filtered CDS were concatenated into super-genes for each species to construct a phylogenetic tree using RAXML v8.2.4 (Stamatakis 2014) with 1,000 ultrafast bootstrap replicates. To estimate divergence times, MCMCtree from the PAML software package was performed on the inferred phylogenetic tree with *D. rerio* as the outgroup and fourfold degenerate sites (4D) extracted from the super-genes. We set four calibration time points (*G. aculeatus–T. rubripes* ~99–127 Ma; *O. niloticus–O. latipes* ~88–139 Ma; *N. melanostomus–P. glenii* ~59-89 Ma; *G. aculeatus–D. rerio* ~206-252 Ma) taken from TimeTree database to calibrate the calculated divergence times.

### Inference of demographic history

We inferred the demographic histories of *P. glenii* and *N. hainanensis* by pairwise sequentially Markovian coalescent (PSMC) analysis (Li and Durbin 2011). NSG short reads used for the genome assemblies were aligned to the two reference genomes using BWA-MEM (Li and Durbin 2009) with default parameters. To generate consensus diploid sequences of the two individuals severally, the SAMtools mpileup with bcftools and vcfutils.pl pipeline (https://github.com/lh3/psmc) was applied. We then used the fq2psmcfa program of PSMC to convert the consensus fastq files into psmcfa format, the input files for PSMC. Finally, the effective population history was inferred using PSMC with 100 bootstraps and plotted by the psmc_plot.pl pileline based on a substitution rate of 2.89e-9 per generation for *P. glenii* and 3.11e-9 per generation for *N. hainanensis*.

### Mutation rate and strength of natural selection

We chose four closely related species for whole-genome synteny alignment using LAST v942 (Kielbasa, et al. 2011) with *O. potamophila* genome sequence used as a reference. The aligned results were submitted to the subprogram “roast” of Multiz v3 (Blanchette, et al. 2004) to generate one-to-one alignment sequences. A sliding window (100kb) along the synteny alignment was applied to estimate the mutation rate. First, the branch lengths for each window were estimated using RAxML (Stamatakis 2014) based on neutral regions (repetitive sequences, regions located within genes and 3kb upstream/downstream) of every window were filtered. Then, the mutation rates were calculated with r8s using the estimated branch lengths and divergence time previously estimated. In addition, we calculated the ω(*ka/ks*) ratios based on 4,550 one-to-one orthologues from the ten teleosts. The free-ratio model, allowing a separate ω for each branch of a tree, from the codeml program in PAML (Yang 2007) was run on the concatenated orthologues and each of the orthologues.

### Gene family expansion and contraction

We used CAFE v3.1(De Bie, et al. 2006) to test for the expansion and contraction of gene families in *P. glenii* based on the results from the OrthoFinder (Emms and Kelly 2015) analyses and the estimated divergence times from MCMCtree. If the copy number of the *P. glenii* was higher or lower than that of its close ancestral branch lineage, then we identified this gene family as substantially expanded or contracted gene family. Any gene family, with a false discovery rate (FDR) adjusted P-value < 0.05, was thought to experience significant expansion or contraction. Functional categories and pathways in significantly expanded gene families were identified by performing GO terms enrichment analysis and KEGG pathway enrichment analysis. GO terms or KEGG pathways with a p-value <0.05 were considered significantly enriched.

### Identification of positively and rapidly evolved genes

All one-to-one orthologous genes were used to assess the contribution of natural selection on the *P. glenii* genome by calculating the ratios (ω) of nonsynonymous substitution (dN) to synonymous substitution (dS) using the software package PAML4 (Yang 2007). The two-ratio branch model (model=2, NSsites=0) was used to detect REGs and branch-site model (model=2, NSsites=2) was used to detect PSGs, with *P. glenii* as the foreground branch. Likelihood ratio tests (LRTs) were applied to test the significance of the differences between alterative and null models for each orthologue. We treated a gene as a REG when the FDR-adjusted p-value < 0.05 and a higher ω ratio in *P. glenii*. Genes with p < 0.05 were considered as PSG. Finally, we removed the false positive results by manual checking. The final REGs and PSGs were then assessed for enrichment of functional categories and pathways.

### Transcriptome sequencing and analysis

Samples were collected from fish in September 2020, late February 2021, and late March 2021 to represent AC, FR and RE stages, respectively, with 6 fish collected at each time point. Brain, muscle, and liver tissue were collected for a total of 54 samples. RNA was extracted from 27 samples (three replicates per stage) using TRIzol (Invitrogen, USA) to generate paired-end (PE) libraries. Each library was sequenced on an Illumina HiSeq platform with 150 bp PE reads. A total of 206.81 Gb clean data were generated (supplementary table S12). Clean reads were mapped to the reference *P. glenii* genome using Hisat2 (Kim, et al. 2015). StringTie (Pertea, et al. 2015) was then used to generate gene expression level counts in fragments per kilobase of transcript per million fragments mapped (FPKM). PCA was performed based on the expression pattern of all genes. To identify DEGs between the different life stages, the number of reads mapped to gene regions was quantified by the FeatureCounts program (Liao, et al. 2014), a part of the Subread package v2.0.0 (http://subread.sourceforge.net/). The R package DESeq2 (Love, et al. 2014) was used for the detection of DEGs based on the read count table generated by featureCounts. An FDR adjusted p value < 0.05 and a 2-fold-change > 2 was set as the level of significance. Enrichment of KEGG pathways and GO terms of the DEGs was estimated using the annotations of all identified transcripts as a background.

### Metabolite extraction, detection, and analysis

Muscle and liver samples from 18 fish (6 replicates per stage) were used for LC–MS/MS analysis. 50mg of each sample was homogenized with 1000 µl of ice-cold methanol/water (70%, v/v). The supernatant was extracted to detect metabolites using a combination of non-targeted detection (Ultra-performance liquid chromatography (UPLC) and Quadrupole-Time of Flight) and widely targeted detection (UPLC and Tandem mass spectrometry (MS/MS)). PCA was performed using statistics function prcomp within R to identify general trends in the content changes. Metabolites with VIP (variable importance in projection) >= 1, absolute Log2FC (fold change) >= 1 and FDR-p value < 0.05 were regarded as SDMs. Identified metabolites were annotated using the KEGG Compound database (http://www.kegg.jp/kegg/compound/), and then annotated metabolites were mapped to the KEGG Pathway database (http://www.kegg.jp/kegg/pathway.html).

### Real-time quantitative PCR assay

To validate DEGs across different life stages of *P. glenii*, qRT-PCR was performed. First strand cDNA was synthesized from 1ug of total RNA samples using Hifair^®^ II 1^st^ Strand cDNA Synthesis SuperMix for qRT-PCR (Yeasen, China). The qRT-PCR was performed with the Hieff^®^ qPCR SYBR^®^ Green Master Mix (Yeasen, China) and a LightCycler 480 II Instrument (Roche, Switzerland). Six biological replicates and three reaction replicates for each group were used. Expression values were calculated using the detected threshold cycle (Ct) value using the geNorm algorithm. Non-tissue-specific reference genes elongation factor alpha (ef1a) and 18S rRNA were selected as internal controls to normalize the relative expression levels. Statistical analysis was performed using an unpaired two-tailed Student’s t test.

## Acknowledgments

This research was supported by the Strategic Priority Research Program of Chinese Academy of Sciences (Grant No. XDB31000000), the National Natural Science Foundation of China (32170480,31972866), Youth Innovation Promotion Association, Chinese Academy of Sciences (http://www.yicas.cn), State Key Laboratory of Genetic Resources and Evolution, Kunming Institute of Zoology, Chinese Academy of Sciences (GREKF21-04), and the Young Top-notch Talent Cultivation Program of Hubei Province. This research was supported by the Wuhan Branch, Supercomputing Center, Chinese Academy of Sciences, China. We also thank Dr Baosheng Wu and Dr Chenguang Feng for their help in our comparative genome analyses.

## Author Contributions

L.Y. and S.H. designed and managed the project. H.J., Y.L., and H.B. collected and prepared the Amur sleeper samples. W.L, Y.Q. and M.M. performed genome assembly and gene annotation. W.L., H.J., and C.W. conducted the bioinformatic analysis and transcriptome, metabolism analysis. W.L., H.J., D.I., N.S., and F.C. wrote and revised the manuscripts.

## Data availability statement

The sequence data have been deposited in the NCBI BioProject database with accession numbers PRJNA818152 (*P. glenii*), PRJNA818180 (*N. hainanensis*). The genome assembly files are under accession numbers JALDQB000000000 (*P. glenii*) and JALDNG000000000 (*N. hainanensis*).

