## supplementary fig. S, supplementary table S, supplementary movie S for "Multiple omic investigations of freeze tolerance adaptation in the aquatic ectothermic vertebrate, the Amur sleeper"

**Supplementary Figures**

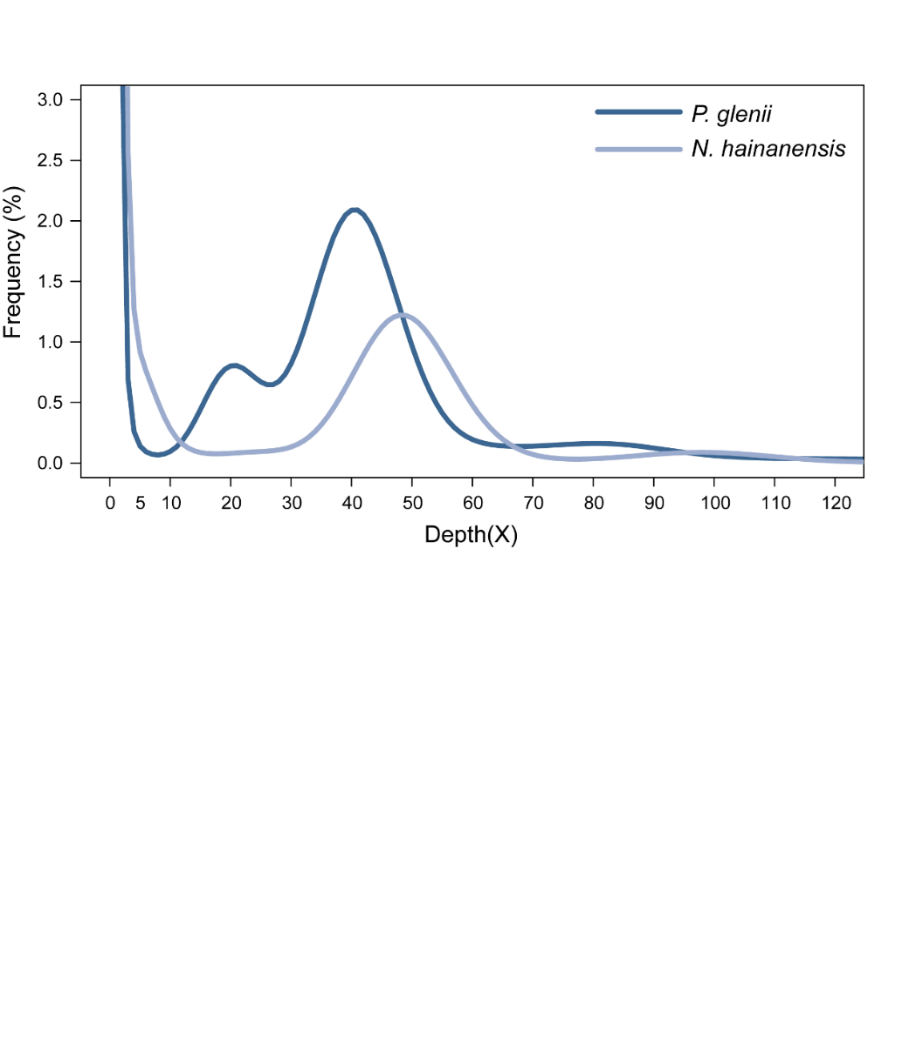

**Supplementary fig. S1.** K-mer (k=17) distribution in *P. glenii* and *N. hainanensis*. The x-axis is depth (X); the y-axis is the proportion which represents the frequency at that depth divided by the total frequency of all the depth.

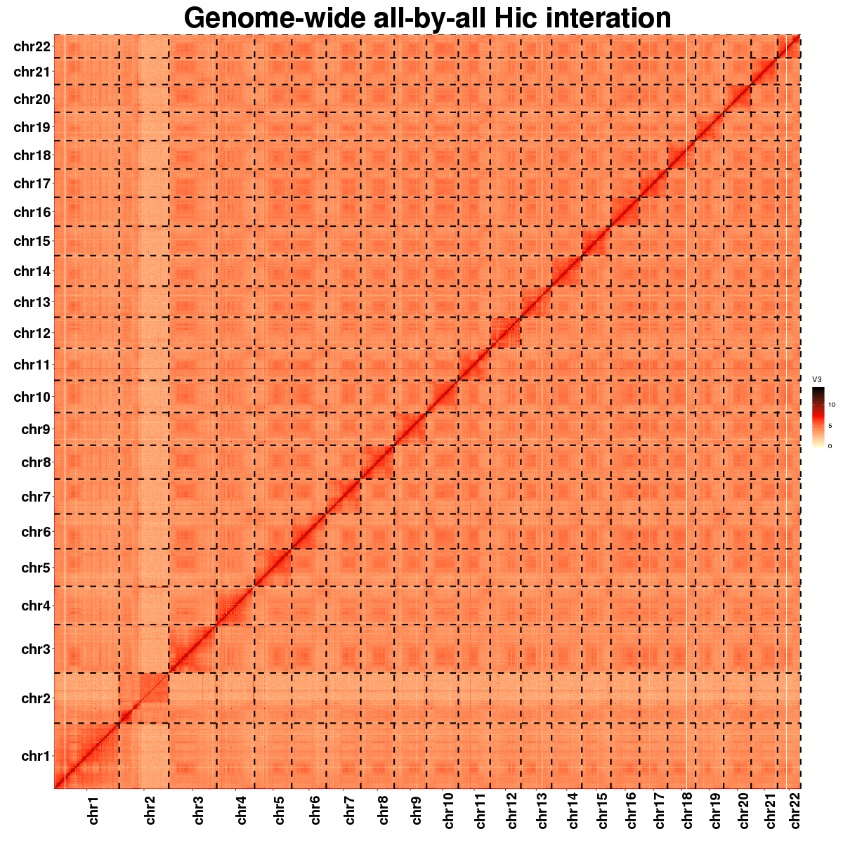

**Supplementary fig. S2.** Genome-wide Hi-C heatmap of *P. glenii*. The blocks represent the 22 pseudochromosomes. The color bar illuminates the contact density from yellow (low) to red (high).

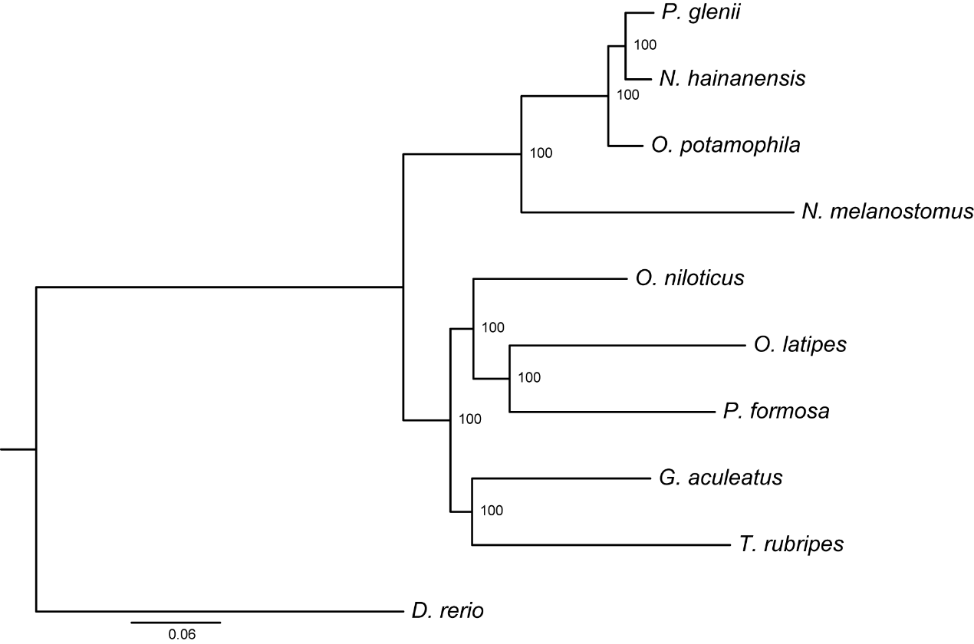

**Supplementary fig. S3.** Phylogenetic tree of *P. glenii* and nine other teleosts. It was constructed from Maximum likelihood methods based on the 4550 concatenated one-to-one orthologues. *D. rerio* was selected as the outgroup.

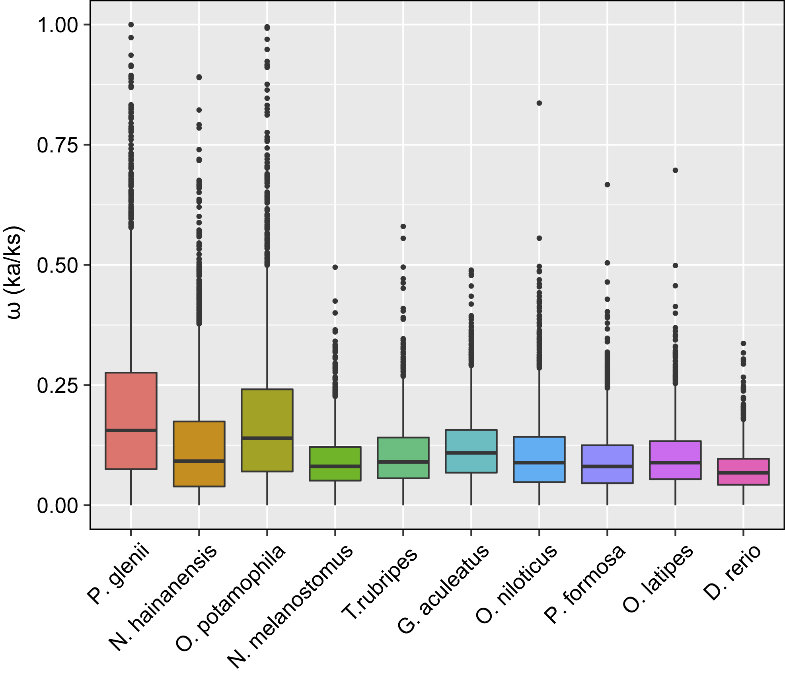

**Supplementary fig. S4.** Boxplot of the Ka/Ks ratios in the ten species. For each species, the black line in the middle indicates the median value. The lower and upper hinges correspond to the first and third quantiles (the 25^th^ and 75^th^ percentiles).

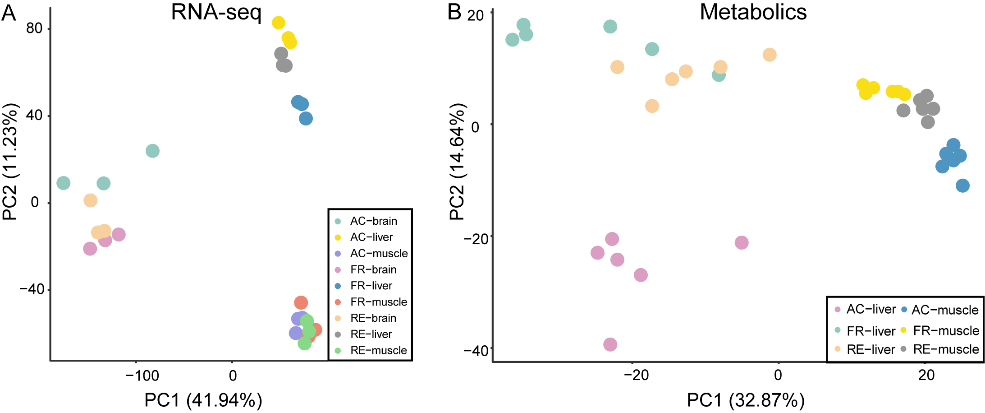

**Supplementary fig. S5.** Principal component analysis (PCA) for RNA-seq (A) and metabolome (B) data during different periods (AC: active autumn; FR: winter freezing; RE: early spring recovery) in three tissues.

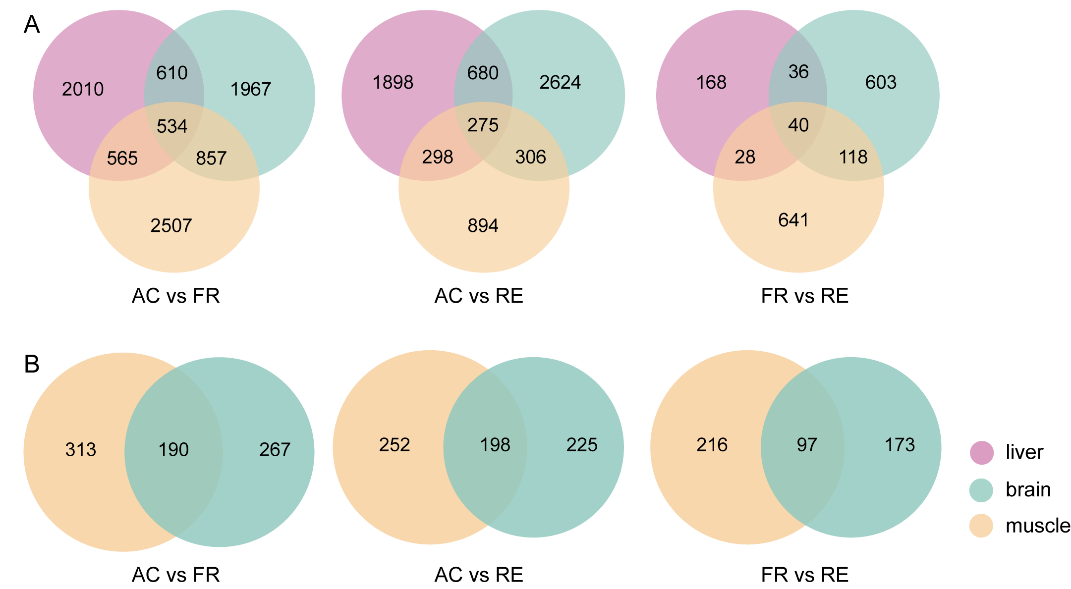

**Supplementary fig. S6.** Venn diagram showing shared and unique variations of gene expression (A) and metabolites (B) in the liver, brain and muscle tissues.

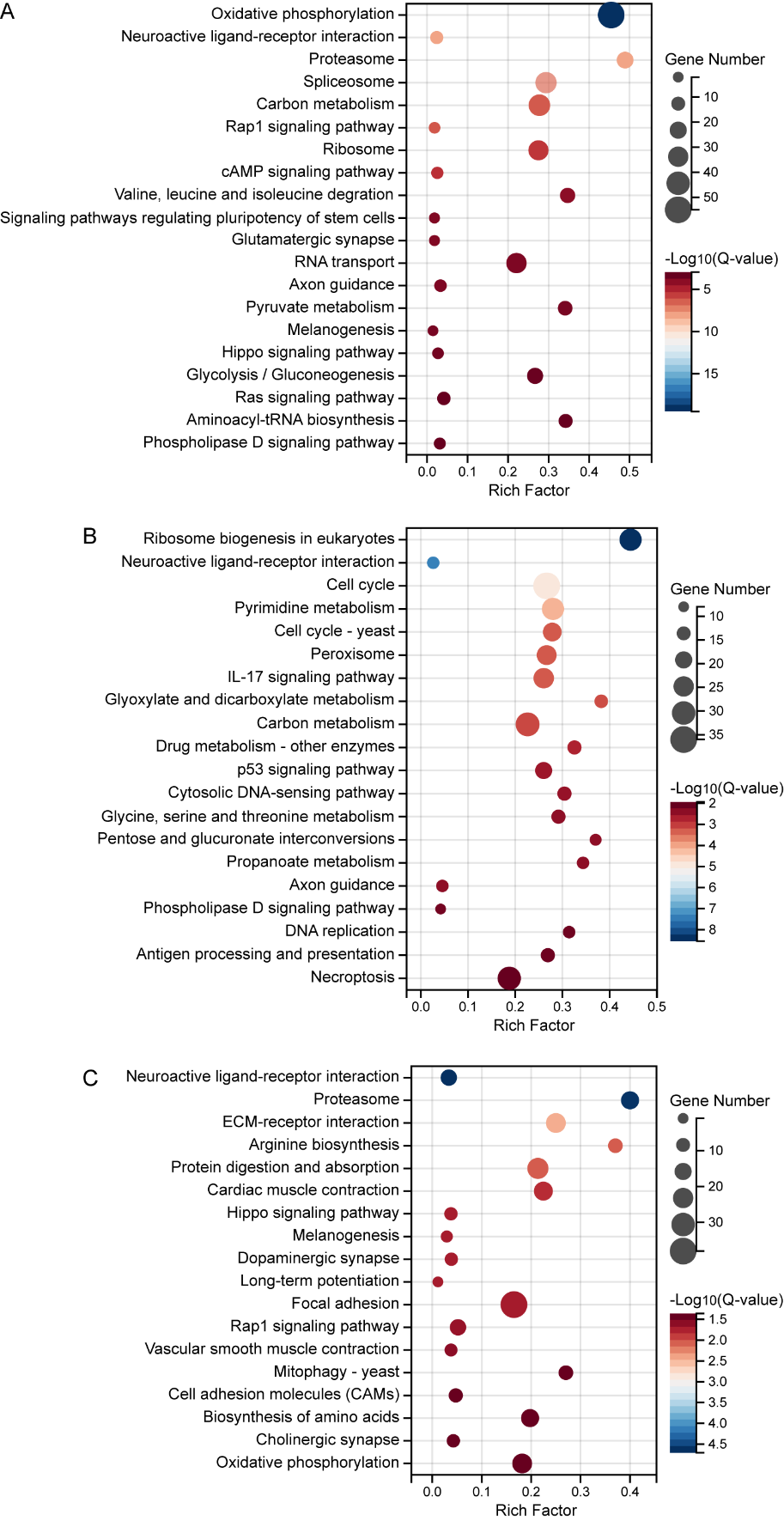

**Supplementary fig. S7.** The kyoto encyclopedia of genes and genomes (KEGG) pathways enrichment of down-regulated genes in the comparison, FR vs AC, in the brain (A), liver (B), and muscle (C) tissues. Only the top 20 categories are shown for brain and liver tissues.

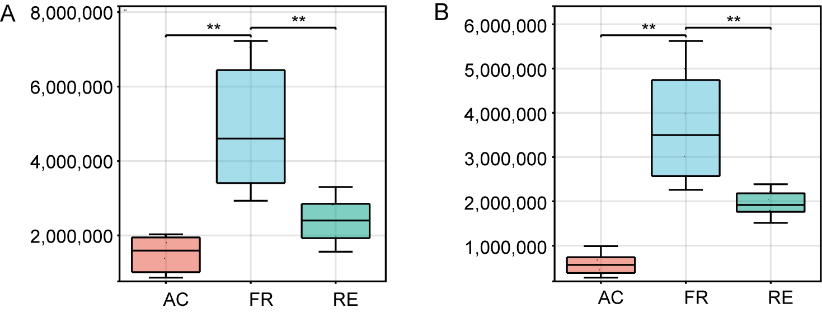

**Supplementary fig. S8.** The relative level of metabolite malonic acid, an inhibitor of tricarboxylic acid cycle, in liver (A) and muscle tissues (B) at three stages (AC: active autumn; FR: winter freezing; RE: early spring recovery). The significance was tested by unpaired two-tailed Student’s t test (**P < 0.01).

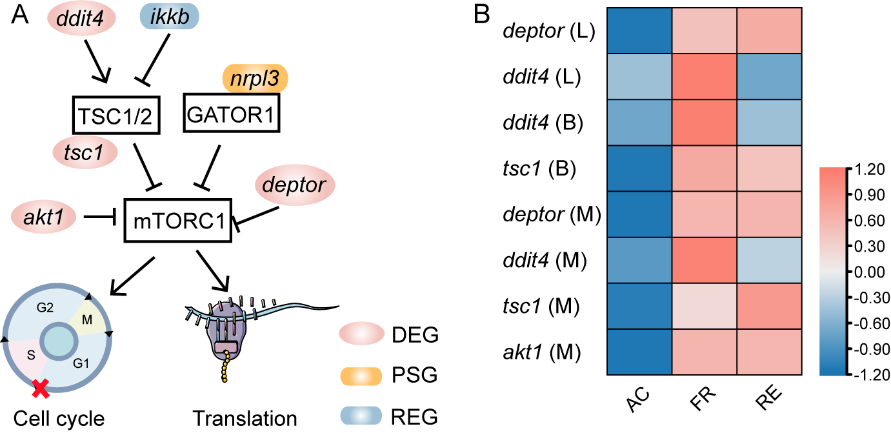

**Supplementary fig. S9.** Schematic depiction of mTORC1 regulation (A) and expression changes of negative regulators in three tissues (L: liver; B: brain; M: muscle). Pink represents higher expression levels, and blue represents lower expression levels.

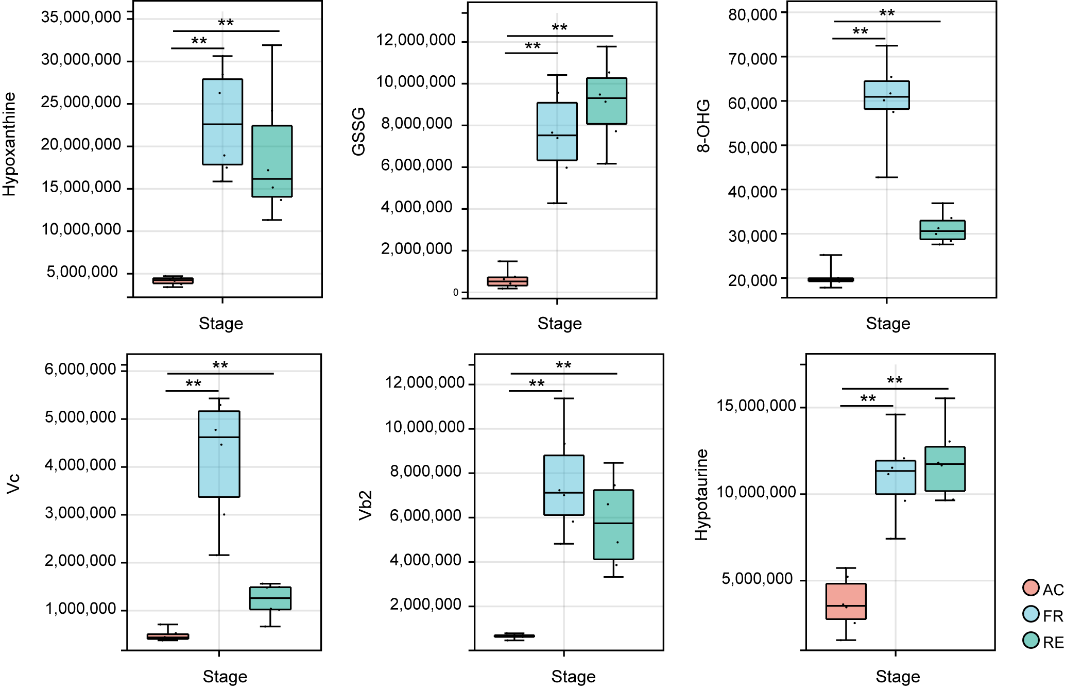

**Supplementary fig. S10.** The relative level of metabolites related to antioxidant defense in muscle tissue during three periods (AC: active autumn; FR: winter freezing; RE: early spring recovery).. The first two metabolites are markers of oxidative stress, while the others are potential antioxidants. Significance was tested by unpaired two-tailed Student’s t test (**P < 0.01).

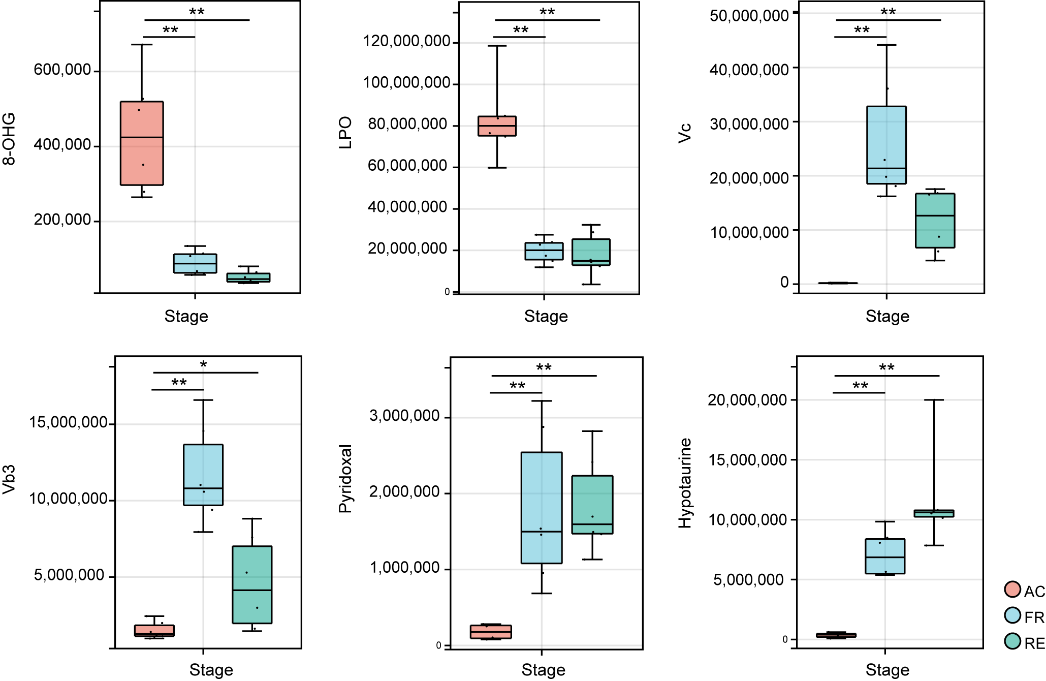

**Supplementary fig. S11.** The relative level of metabolites related to antioxidant defense in liver tissue during three periods (AC: active autumn; FR: winter freezing; RE: early spring recovery).. The first two metabolites are markers of oxidative stress, while the others are potential antioxidants. Significance was tested by unpaired two-tailed Student’s t test (*P < 0.05; **P < 0.01).

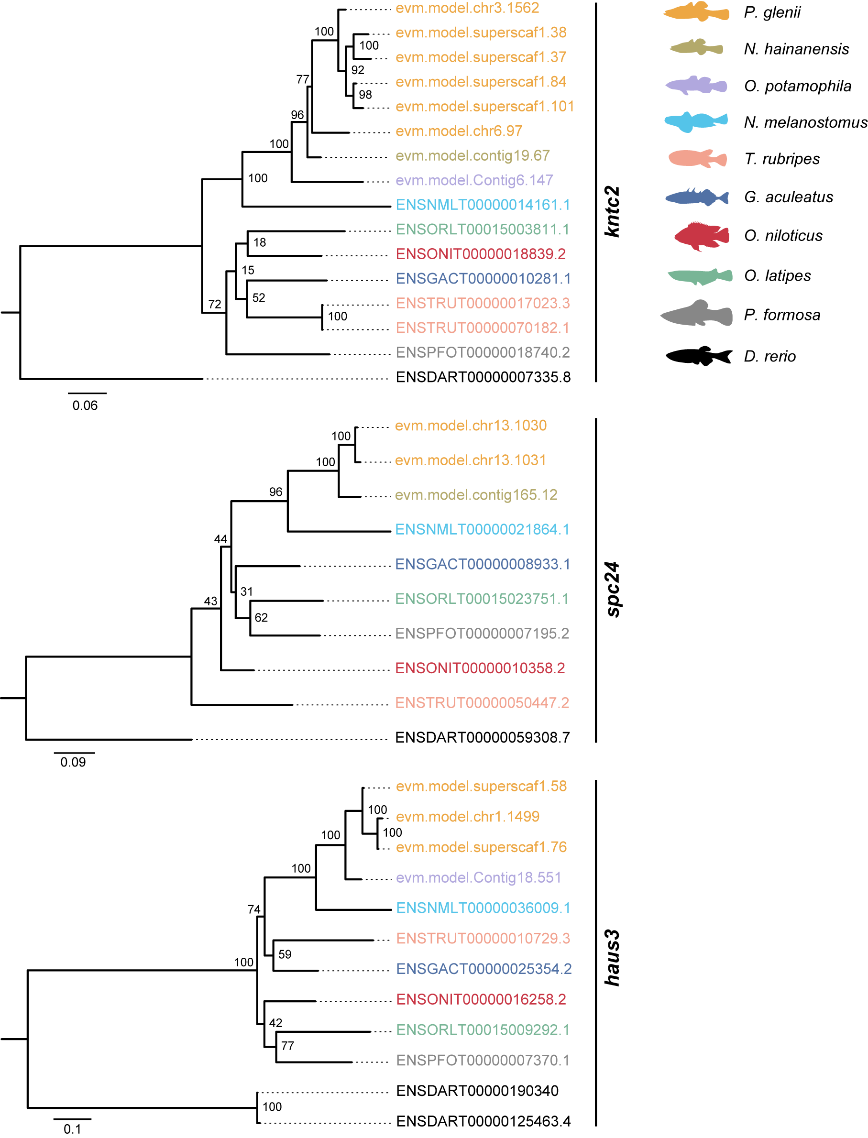

**Supplementary fig. S12.** The phylogenetic analysis of expanded genes involved in cytoskeleton using maximum likelihood methods. The number on the node of each branch is the bootstrap value.

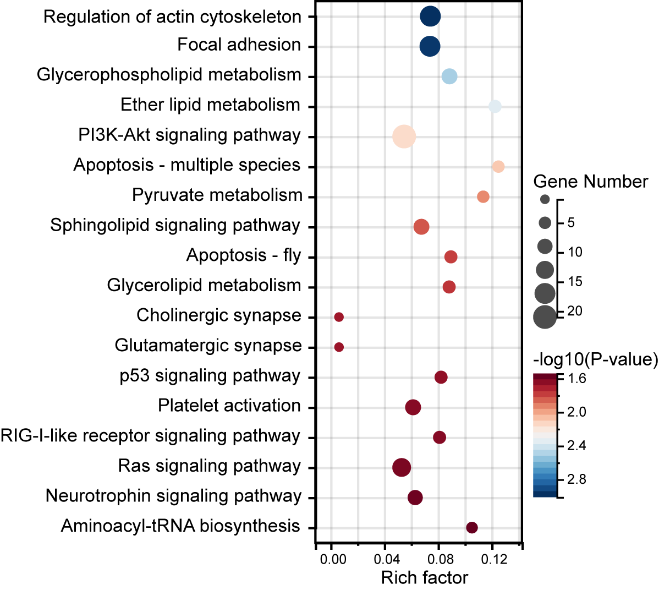

**Supplementary fig. S13.** The kyoto encyclopedia of genes and genomes (KEGG) pathways enrichment of rapidly evolving genes. Regulation of actin cytoskeleton was mostly significant enriched.

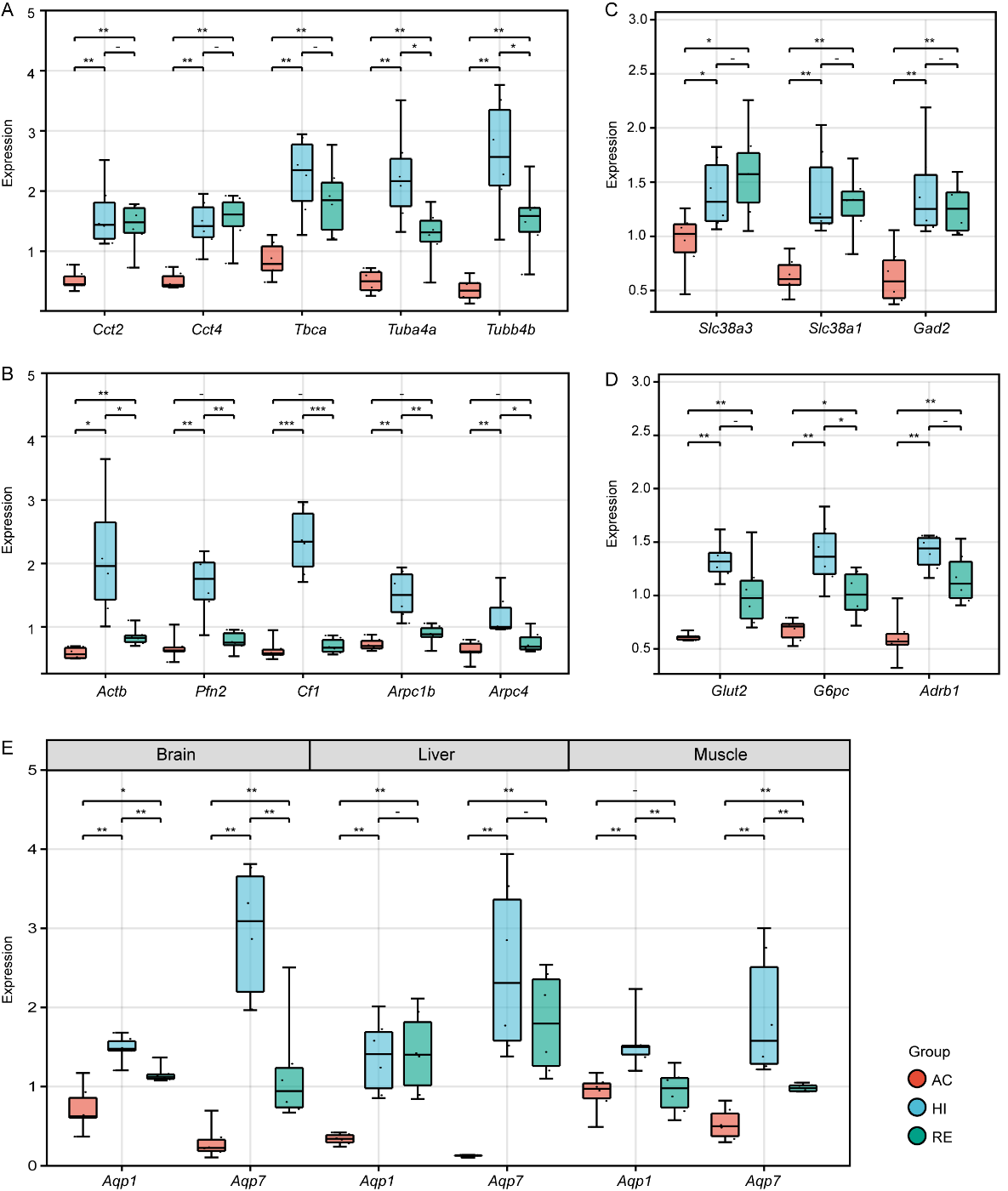

**Supplementary fig. S14.** Expression levels of differentially expressed genes (DEGs) picked randomly from results of transcriptome analyses. (A) Expression of DEGs associated with microtubule in muscle tissue during three stages. (B) Expression of DEGs associated with actin cytoskeleton in muscle tissue during three stages. (C) Expression of DEGs in GABAergic synaptic transmission of brain tissue. (D) Expression of DEGs associated with glucose accumulation in liver tissue. (E) Expression of aquaporins in three tissues. Each stage included 6 biological replicates (n=6). The significance was tested by unpaired two-tailed Student’s t test (*P < 0.05; **P < 0.01; NS-not significant, P > 0.05).

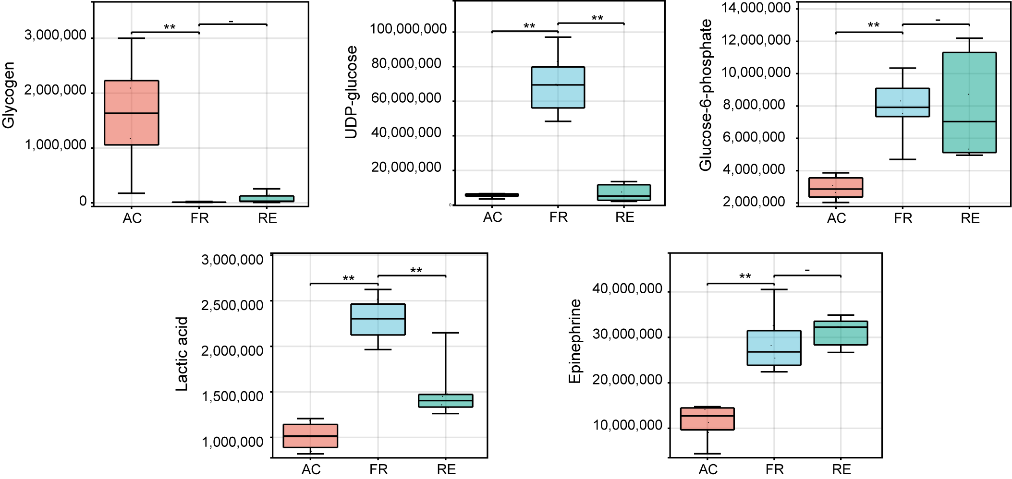

**Supplementary fig. S15.** The relative level of metabolites involved in glycogenolysis and glycolysis in liver tissue during three periods (AC: active; FR: freezing; RE: recovery; L: liver; B: brain; M: muscle). The significance was tested by unpaired two-tailed Student’s t test (**P < 0.01; NS-not significant, P > 0.05).

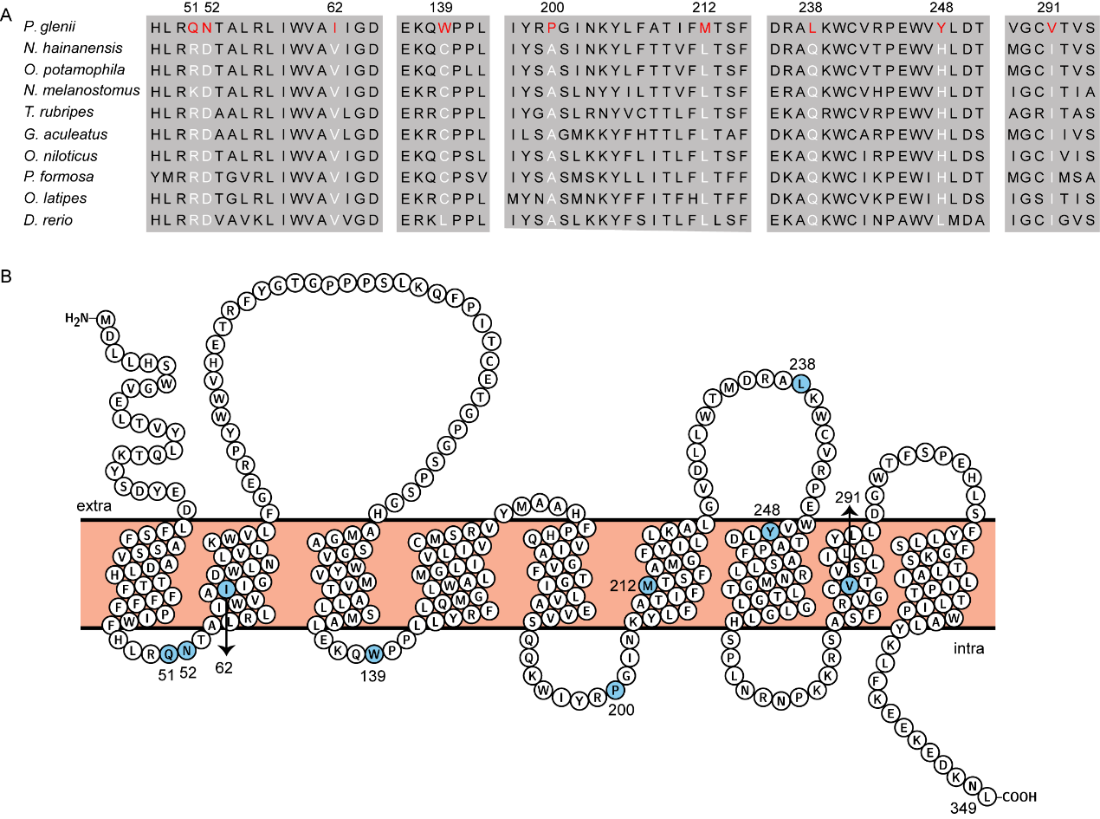

**Supplementary fig. S16.** Specific mutations in glucose-6-phosphatase (*g6pc*). (A) The sequence alignment of *g6pc* of the ten teleosts with specific mutations in Amur sleeper marked by red. (B) 2D transmembrane protein diagram of g6pc was shown. The specific mutations have blue background.

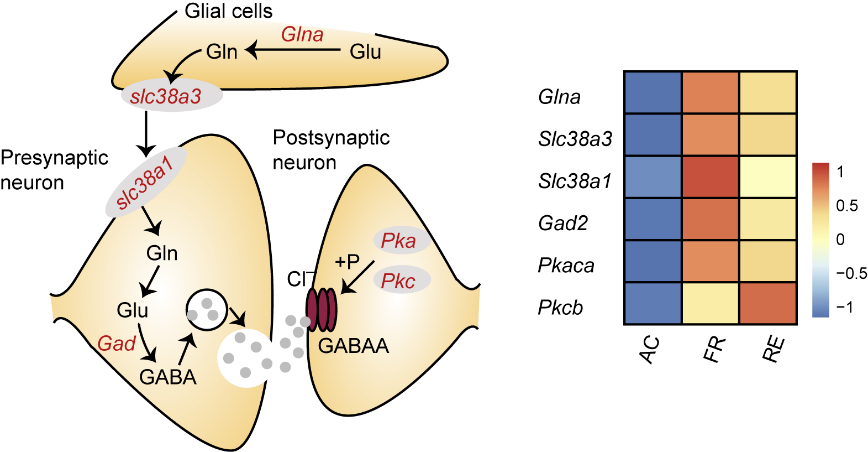

**Supplementary fig. S17.** The schematic of GABAergic synapse, and expression changes of genes related to the pathway during three periods (AC: active autumn; FR: winter freezing; RE: early spring recovery) in the brain tissue. Dark orange represents higher expression levels, and dark blue represents lower expression levels.

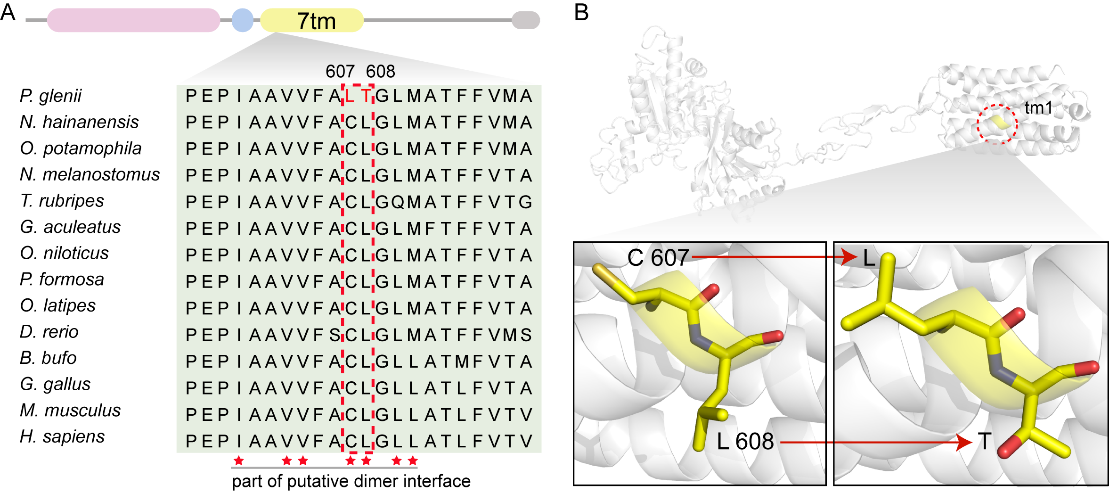

**Supplementary fig. S18.** Positively selected sites in metabotropic glutamate receptor 5 (*mglur5*). (A) Sequence alignment of adjacent region of positive selection sites, and the mutations in Amur sleeper were marked with red. The sites pointed with red pentagram form part of putative dimer interface that is inferred by searching for conserved domains in NCBI. (B) Three-dimensional (3D) structure of the Amur sleeper mglur5 simulated by homologous approach based on the 3D structure of human mglur5 (PDB ID: 7FD9), was shown, the two positively selected sites were marked by yellow.

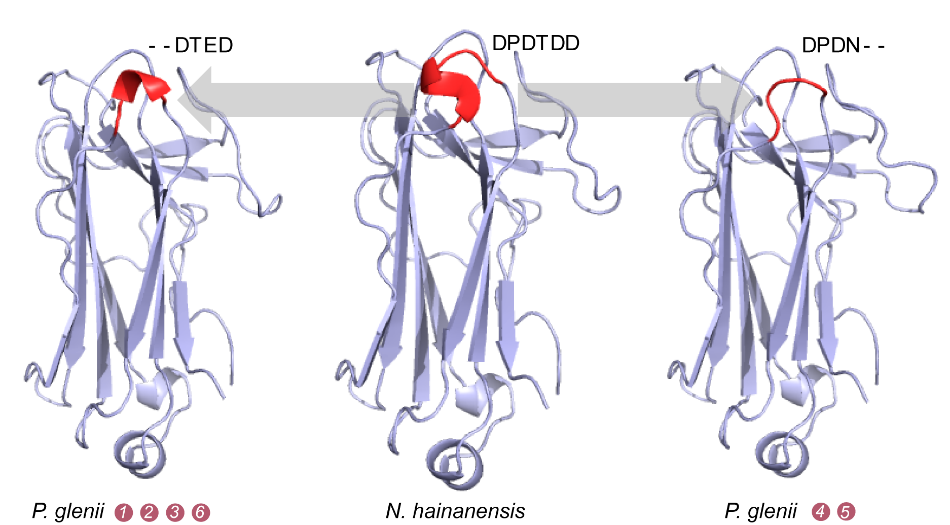

**Supplementary fig. S19.** Three-dimensional (3D) structure of the Amur sleeper f-box only protein 2 (fbox2) simulated by homologous approach with the 3D structure of mouse fbox2 (PDB ID: 1UMI) as model. The structures marked by red represent six continuous amino acid (from the 122^nd^ to the 127^th^). The deletion of two-amino acids caused the structural changes of Amur sleeper fbox2.

**Supplementary Tables**

**Supplementary table S1. Sequence information used in genome assembly.**

| Sequencing type | *P. glenii* | *N. hainanensis* |
| --- | --- | --- |
| Nanopore sequencing (Gb) | 70.29 | 86.06 |
| BGI sequencing (Gb) | 101.02 | 121.71 |
| Hic sequencing (Gb) | 95.60 | - |
| Total (Gb) | 266.91 | 207.77 |

**Supplementary table S2.** Estimation of genome size based on 17-mer statistics.

|  | Kmer | Depth | N-kmer | Genome size (Mb) | Heterozygous rate (%) |
| --- | --- | --- | --- | --- | --- |
| *P. glenii* | 17 | 41.12 | 34,413,408,900 | 827.25 | 0.64 |
| *N. hainanensis* | 17 | 49.04 | 42,682,290,335 | 840.86 | 0.15 |

**Supplementary table S3. Comparison of *P. glenii* genome assembly using Wtdbg, Flye and Smartdenovo.**

| State type | Wtdbg | Flye | Smartdenovo |
| --- | --- | --- | --- |
| Genome size (Mb) | 762.19 | 700.22 | 710.22 |
| Longest contig (Mb) | 22.08 | 13.91 | 21.27 |
| Contig N50 (Mb) | 1.46 | 0.97 | 5.49 |
| Contig N90 (Kb) | 65.53 | 165.21 | 550.20 |

**Supplementary table S4.** Statistics of assembled 22 chromosomes of ***P. glenii*** genome.

|  | Length(bp) | % of genome | Contig number |
| --- | --- | --- | --- |
| Chr1 | 59,894,370 | 0.0843 | 38 |
| Chr2 | 45,882,592 | 0.0646 | 298 |
| Chr3 | 44,353,964 | 0.0624 | 34 |
| Chr4 | 34,837,245 | 0.0490 | 39 |
| Chr5 | 34,319,016 | 0.0483 | 19 |
| Chr6 | 31,962,909 | 0.0450 | 14 |
| Chr7 | 31,895,894 | 0.0449 | 26 |
| Chr8 | 30,990,602 | 0.0436 | 23 |
| Chr9 | 30,140,547 | 0.0424 | 20 |
| Chr10 | 29,556,609 | 0.0416 | 23 |
| Chr11 | 29,157,000 | 0.0410 | 21 |
| Chr12 | 28,553,870 | 0.0402 | 19 |
| Chr13 | 28,359,885 | 0.0399 | 28 |
| Chr14 | 27,802,162 | 0.0391 | 11 |
| Chr15 | 26,947,343 | 0.0379 | 22 |
| Chr16 | 26,416,522 | 0.0372 | 13 |
| Chr17 | 26,031,007 | 0.0366 | 15 |
| Chr18 | 25,779,153 | 0.0363 | 20 |
| Chr19 | 25,751,100 | 0.0363 | 22 |
| Chr20 | 25,544,473 | 0.0360 | 9 |
| Chr21 | 24,206,277 | 0.0340 | 15 |
| Chr22 | 21,250,410 | 0.0299 | 16 |
| Total | 689,632,950 | 0.9709 | 702 |

**Supplementary table S5. Genome assembly statistics for *P. glenii* and *N. hainanensis*.**

| State type | *P. glenii* | | *N. hainanensis* |
| --- | --- | --- | --- |
|  | Contig | Scaffolds | Contig |
| Number | 2213 | 1427 | 8221 |
| Total bases (bp) | 710,217,920 | 710,296,520 | 848,143,664 |
| Chromosome number | **-** | 22 | **-** |
| N50 (bp) | 2,962,197 | 29,556,609 | 1,344,149 |
| N90 (bp) | 348,803 | 25,544,473 | 37,215 |
| Maximum (bp) | 12,999,775 | 59,894,370 | 22,209,510 |
| Minimum (bp) | 48 | 48 | 2,600 |
| Average length (bp) | 320,929 | 497,755 | 103,167 |
| GC content (%) | 39.80 | 39.80 | 39.38 |
| Total N | 0 | 78,600 | 0 |
| %N | 0 | 0.0001 | 0 |

**Supplementary table S6. Short reads coverage on two assembly genomes.**

|  |  | *P. glenii* | *N. hainanensis* |
| --- | --- | --- | --- |
| Reads | Number of clean reads | 1,010,068,256 | 811,389,940 |
|  | Percentage of mapped reads | 99.42 | 97.21 |
| Genome | Coverage（%） | 98.61 | 97.16 |
|  | Coverage at least 5×（%） | 97.72 | 95.34 |
|  | Coverage at least 10×（%） | 97.03 | 93.81 |

**Supplementary table S7. BUSCO assessment results of *P. glenii* and *N. hainanensis*.**

|  | *P. glenii* | *N. hainanensis* |
| --- | --- | --- |
| Complete BUSCOs (C) | 4,182 (91.2%) | 4,268 (93.1%) |
| Complete and single-copy BUSCOs (S) | 4,068 (88.7%) | 4,101 (89.5%) |
| Complete and duplicated BUSCOs (D) | 114 (2.5%) | 167 (3.6%) |
| Fragmented BUSCOs (F) | 150 (3.3%) | 124 (2.7%) |
| Missing BUSCOs (M) | 252 (5.5%) | 192 (4.2%) |
| Total Lineage BUSCOs | 4,584 | 4,584 |

**Supplementary table S8. Statistics of repetitive sequences in *P. glenii* and *N. hainanensis.***

|  |  | Repbase TEs | | TE proteins | | *De novo* | | Combined TEs | |
| --- | --- | --- | --- | --- | --- | --- | --- | --- | --- |
| Species | Type | Lengths (bp) | Percent in genome (%) | Lengths (bp) | Percent in Genome (%) | Lengths (bp) | Percent in genome (%) | Lengths (bp) | Percent in genome (%) |
| *P. glenii* | DNA | 14101793 | 1.74 | 10209910 | 1.26 | 127666508 | 15.72 | 130308576 | 16.05 |
|  | LINE | 32592950 | 4.01 | 44320284 | 5046 | 72083804 | 8.88 | 81890017 | 10.08 |
|  | SINE | 7286574 | 0.90 | 0 | 0.00 | 13041380 | 1.61 | 14795810 | 1.82 |
|  | LTR | 4662699 | 0.57 | 14765931 | 1.82 | 38080511 | 4.69 | 43132382 | 5.31 |
|  | Satelite | 3236110 | 0.40 | 0 | 0.00 | 5209407 | 0.64 | 8388799 | 1.03 |
|  | SSR | 11256663 | 1.39 | 0 | 0.00 | 4498174 | 0.55 | 57885642 | 7.13 |
|  | Unknow | 0 | 0.00 | 0 | 0.00 | 53647128 | 6.61 | 53647128 | 6.61 |
|  | Total | 73263864 | 9.02 | 69296125 | 8.53 | 314884874 | 38.78 | 390833391 | 48.13 |
| *N. hainanensis* | DNA | 7641811 | 0.90 | 7695660 | 0.91 | 78888418 | 9.30 | 82790660 | 9.76 |
|  | LINE | 26042472 | 3.07 | 36192234 | 4.27 | 61168715 | 7.21 | 66812219 | 7.88 |
|  | SINE | 5672350 | 0.67 | 0 | 0.00 | 10740124 | 1.27 | 12483154 | 1.47 |
|  | LTR | 4028390 | 0.47 | 19161553 | 2.26 | 67009458 | 7.90 | 73655552 | 8.68 |
|  | Satellite | 814992 | 0.10 | 0 | 0.00 | 3711606 | 0.44 | 4107392 | 0.48 |
|  | SSR | 8802658 | 1.04 | 0 | 0.00 | 19232498 | 2.27 | 89554670 | 10.56 |
|  | Unknow | 0 | 0.00 | 0 | 0.00 | 59781152 | 7.05 | 59781152 | 7.05 |
|  | Total | 53135699 | 6.26 | 63049447 | 7.43 | 301269110 | 35.52 | 390054964 | 45.99 |

**Supplementary table S9. Summary statistics of gene function annotation.**

|  | | *P. glenii* | | *N. hainanensis* | |
| --- | --- | --- | --- | --- | --- |
|  | | number | percent (%) | number | percent (%) |
| Annotated | InterPro | 21,628 | 91.71 | 22,133 | 84.36 |
|  | GO | 18,757 | 79.54 | 20,204 | 77.01 |
|  | KEGG | 14,908 | 63.22 | 12,829 | 48.90 |
|  | Swissprot | 20,377 | 86.41 | 21,903 | 83.48 |
|  | Nr | 22,104 | 93.73 | 23,997 | 91.46 |
| Number of annotated genes | | 22,762 | 96.52 | 24,827 | 94.63 |
| Number of unannotated genes | | 820 | 3.48 | 1410 | 5.37 |

**Supplementary table S10.** **Sequence information of RNA-seq used in genome annotation.**

|  | *P. glenii* | | | *N. hainanensis* | | |
| --- | --- | --- | --- | --- | --- | --- |
| Tissue | Clean reads | Clean base (Gp) | Q20(%) | Clean reads | Clean base (bp) | Q20(%) |
| Muscle | 41,814,802 | 6.27 | 97.07 | 63,766,838 | 9.57 | 96.77 |
| Skin | 42,309,194 | 6.27 | 97.07 | 64,666,720 | 9.707 | 96.81 |
| Heart | 40,814,802 | 6.13 | 97.22 | 60,670,798 | 9.107 | 97.58 |
| Liver | 41,43,094 | 6.22 | 97.15 | 65,395,876 | 9.81 | 96.92 |
| Spleen | 41,991,306 | 6.30 | 97.20 | 62,170,602 | 9.33 | 96.87 |
| Brain | 41,749,126 | 6.26 | 97.18 | 67,603,410 | 10.14 | 97.35 |
| Eye | 41,371,728 | 6.26 | 97.14 | 62,363,824 | 9.35 | 96.89 |
| Gonad | 40,991,754 | 6.15 | 97.18 | 66,697,846 | 10.00 | 96.97 |
| Kidney | 41,746,764 | 6.15 | 97.32 | - | - | - |
| Blood | 42,247,430 | 6.34 | 97.00 | - | - | - |
| Gill | - | - | - | 68,658,810 | 10.29 | 96.86 |
| Total | 375,036,906 | 62.49 | - | 581,994,724 | 87.30 | - |

**Supplementary table S11.** **Sequence information of full-length transcriptome used in genome annotation.**

| Species | Subreads | Total length  (bp) | Average length (bp) | Maximal length  (bp) |
| --- | --- | --- | --- | --- |
| *P. glenii* | 8,570,679 | 17,223,271,319 | 2,009 | 113,774 |
| *N. hainanensis* | 16,621,922 | 28,577,594,640 | 1,179 | 186,637 |

**Supplementary table S12.** **Sequence information for RNA-seq of Amur sleeper in different periods.**

| Stage | Sample | Clean Reads | Clean Base(Gb) | Q30(%) |
| --- | --- | --- | --- | --- |
| Active | Brain-1 | 50,255,350 | 7.54 | 92.94 |
|  | Brain-2 | 41,050,656 | 6.16 | 93.25 |
|  | Brain-3 | 46,754,146 | 7.01 | 93.22 |
|  | Liver-1 | 42,207,234 | 6.33 | 93.95 |
|  | Liver-2 | 46,760,598 | 7.01 | 93.69 |
|  | Liver-3 | 41,546,454 | 6.23 | 93.62 |
|  | Muscle-1 | 45,848,112 | 6.88 | 93.39 |
|  | Muscle-2 | 47,348,984 | 7.1 | 93.27 |
|  | Muscle-3 | 50,230,586 | 7.53 | 93.19 |
| Freezing | Brain-1 | 54,532,896 | 8.18 | 94.57 |
|  | Brain-2 | 53,211,314 | 7.98 | 94.43 |
|  | Brain-3 | 52,026,806 | 7.8 | 94.14 |
|  | Liver-1 | 53,504,234 | 8.03 | 94.75 |
|  | Liver-2 | 55,192,844 | 8.28 | 94.29 |
|  | Liver-3 | 46,636,744 | 7 | 93.96 |
|  | Muscle-1 | 43,270,292 | 6.49 | 92.66 |
|  | Muscle-2 | 69,953,852 | 10.49 | 91.89 |
|  | Muscle-3 | 69,277,858 | 10.39 | 91.85 |
| Recovery | Brain-1 | 55,531,632 | 8.33 | 93.79 |
|  | Brain-2 | 46,518,192 | 6.98 | 94.56 |
|  | Brain-3 | 56,708,724 | 8.51 | 94.06 |
|  | Liver-1 | 42,823,136 | 6.42 | 95.11 |
|  | Liver-2 | 45,935,058 | 6.89 | 94.57 |
|  | Liver-3 | 51,780,442 | 7.77 | 94.07 |
|  | Muscle-1 | 61,776,882 | 9.27 | 94.56 |
|  | Muscle-2 | 53,323,192 | 8 | 94.67 |
|  | Muscle-3 | 54,740,670 | 8.21 | 94.56 |
| Total | - | 1,378,746,888 | 206.81 | - |

**Supplementary Movies**

**Supplementary movie S1.** The video showed the Amur sleeper during winter freezing with the entire body frozen in ice.

**Supplementary movie S2.** The video showed the Amur sleeper during early spring recovery.
